# Non-essential kinetochore proteins contribute to meiotic chromosome condensation through polo-like kinase

**DOI:** 10.1101/2023.10.24.563891

**Authors:** Deepika Trakroo, Prakhar Agarwal, Anushka Alekar, Santanu Kumar Ghosh

## Abstract

Chromosome condensation plays a pivotal role during faithful chromosome segregation, hence understanding the factors that drive condensation is crucial to get mechanistic insight into chromosome segregation. Previously we showed that in budding yeast, the absence of the non-essential kinetochore proteins affects chromatin-condensin association in meiosis but not in mitosis. A differential organization of the kinetochores, that we and others observed earlier during mitosis and meiosis may contribute to the meiotic-specific role. Here, with our in-depth investigation using in vivo chromosome condensation assays in cells sans a non-essential kinetochore protein, Ctf19, we establish that these proteins have roles in achieving a higher meiotic condensation without influencing much of the mitotic condensation. We further observed an accumulation of the polo-like kinase Cdc5 owing to its higher protein stability in *ctf19Δ* meiotic cells. High Cdc5 activity causes hyper-phosphorylation of the condensin resulting in its reduced stability and concomitant decreased association with the chromatin. Overall, our findings highlight the role of Ctf19 in promoting meiotic chromosome condensation by influencing the activity of Cdc5 and thereby affecting the stability and association of condensin with the chromatin.

## Introduction

Faithful chromosome segregation across eukaryotes requires that the chromosomes become condensed and attached properly to the microtubule spindle during cell division. The attachment is mediated by a conserved multiprotein complex called ‘kinetochore’ formed at the centromeres of the chromosomes (Cleveland et al., 2003, Ishi and Akiyoshi., 2022). The proteins within a kinetochore are organized in a hierarchical fashion with respect to the centromeric DNA and the microtubules (Biggins., 2013, Cheeseman and Desai., 2008, De Wulf et al., 2003, Westermann et al., 2007). The inner and the outer proteins associate with the centromeres and microtubules, respectively whereas the central proteins act as linker (Meraldi et al., 2006, Dong et al., 2022, Dou et al., 2023). Over the last three decades studies on kinetochore from myriad organisms have revealed that this protein complex, besides establishing a connection with the microtubules, also controls other cellular activities that facilitate high-fidelity chromosome segregation (Glynn et al., 2004, Muller et al., 2012, Meyer et al., 2015, Kim et al., 2013). In budding and fission yeasts, the kinetochore proteins, in particular the essential inner and central proteins, are also shown to promote chromosome condensation (Kruitwagen et al., 2018, Nakazawa et al., 2008).

Chromosome condensation is promoted by an evolutionarily conserved condensin complex, which binds to the chromatin facilitating its condensation through DNA compaction, formation of DNA loops, and its axial shortening (Freeman et al., 2000, Strunnikov et al., 1995, Kruitwagen et al., 2015). Condensation in budding yeast is established in prophase and persists till the end of mitotic division (Vas et al., 2007, Lamothe et al., 2020). The cell cycle-dependent chromatin-condensin association is regulated by several factors. While CDKs initiate this during start of the cell cycle, Polo-like kinase and Aurora B kinase maintain it during metaphase, possibly through phosphorylation of their substrates, including the condensin complex (Giet et al., 2001, Lavoie et al., 2004, St. Pierre et al., 2009). Such timely association and resulting accurate folding of the chromatin are critical for faithful chromosome segregation.

The kinetochore complex also plays a crucial role in chromatin-condensin association.. Several studies have revealed that this role is executed through kinetochore-mediated centromeric and pericentromeric recruitment of several proteins including cohesins (Guacci et al., 1997, Lavoie et al., 2004), shugoshin (Peplowska et al., 2014, Kruitwagen et al., 2018, Yahya et al., 2020), 14-3-3 protein, Bmh1 (Jain et al., 2021) and enzymes such as aurora B kinase, Ipl1 (Giet et al., 2001, Petersen et al., 2003, Lavoie et al., 2004,), polo-like kinase, Cdc5 (St. Pierre et al., 2009, Bazile et al., 2010, Leonard et al., 2015, Lamothe et al., 2020) and a histone deacetylase, Hst2 (Wilkins et al., 2014, Kruitwagen et al., 2015, 2018, Jain et al., 2021). However, the non-essential proteins of the Ctf19 sub-complex (Ctf19c) of the kinetochore (Mehta et al., 2022) were not shown to hinder chromosome condensation (Kruitwagen et al., 2018) as expected from their dispensability in mitotic growth. However, we have shown earlier that in absence of these non-essential proteins, meiosis is severely affected as several meiosis specific events are perturbed (Ghosh et al., 2004, Mehta et al., 2014, Agarwal et al., 2015). This happens as there appears to be a differential organization of the kinetochore assembly between mitosis and meiosis (Mehta et al., 2014, Borek et al., 2021) for which the non-essential proteins perhaps facilitate certain protein-protein and/or protein-DNA interactions in meiosis but not in mitosis.

Strikingly, earlier we observed that Ctf19 has a role to play in meiotic chromosome condensation since the cells in the absence of Ctf19 harbor defects in rDNA condensation and show reduced association of condensin with the chromatin in meiosis but not in mitosis (Mehta et al., 2014). While why chromatin-condensin association is compromised in *ctf19Δ* meiotic cells is not known, we hypothesize that to facilitate prophase I events, meiosis may require extra chromosome condensation than mitosis, and the nonessential kinetochore proteins might contribute to regulating that. This notion is also fueled by the distinct thread-like appearance of the paired homologous chromosomes in pachytene of meiosis I, which is never visible in mitosis and unlikely to happen in meiosis solely due to pairing.

In this report, to test the above hypothesis we sought to compare the extent of chromosome condensation between mitosis and meiosis and to find a molecular explanation of condensation defect in the non-essential kinetochore mutants. Interestingly, we observed using three cell biological in vivo assays that the chromosomes indeed appeared more condensed in meiosis than in mitosis. We further demonstrate that lack of Ctf19 causes misregulation of Cdc5 specifically in meiosis, which leads to hyper-phosphorylation of condensin resulting its dissociation from the chromatin and defect in chromosome condensation. Additionally, using wild type cells overexpressing Cdc5, we show that the misregulation of Cdc5 is indeed the reason for condensin failing to associate properly with the chromatin. This study highlights yet another meiosis-specific essential contribution of the mitotically dispensable kinetochore proteins.

## Results

### Chromosomes are more condensed in meiosis as compared to mitosis

We hypothesize that the level and determinants of chromosome condensation might differ between mitosis and meiosis because earlier, using Ctf19c mutant, we observed that the kinetochore has a role in rDNA condensation in meiosis but not in mitosis (Mehta et al., 2014). However, whether such a role is confined to a specific condensed locus like rDNA or it is observed at other chromosomal locales remains unknown. To investigate this, we first sought out to compare the extent of chromosomal condensation of the kinetochore proximal pericentromeric regions between mitosis and meiosis. Diploid strains were constructed where one copy of chromosome V was marked with [TetO]_224_ –[TetR-GFP] system at a locus 1.4 kb away from the *CEN V* (henceforth called *CEN V*-GFP). The fluorescence intensity of *CEN V*-GFP was used as a surrogate marker to decipher the in vivo condensation level, as described before (Kruitwagen et al., 2015). In this microscopy-based assay, the fluorescence intensity of TetR-GFP bound to the [TetO]_n_ array inserted at a decondensed locus will be more compared to when the array is inserted at a condensed locus. This is because, due to condensation, a linear shrinkage of the operators happens, which brings the TetR-GFP molecules closer to each other, causing quenching of the fluorescence and, hence, reduction in the intensity (Fig. 1A). The *CEN V*-GFP marked strains also harbored SPBs (Spc42) marked with mCherry to judge different cell cycle stages. The GFP signal intensities were measured in the cells residing at the pre-S phase (interphase) stage or at metaphase/metaphase I, which is presumed to contain decondensed and condensed chromatin, respectively. An unbudded cell with a single SPB was taken as a pre-S phase cell in mitosis or meiosis; a large budded cell with two separated SPBs in the mother compartment was taken as a metaphase cell (Fig. 1B) whereas same in an unbudded meiotic cell was considered as a metaphase I cell (Fig. 1C). The timepoint to harvest meiotic pre-S phase cells was determined based on Ime1 (an inducer of meiosis in yeast) expression and FACS analysis. The Ime1 expression is induced by an acetate-based rich medium (YPA), however, it starts accumulating in the nucleus after the depletion of G1 cyclins (Colomina et al., 1999, 2003). In our experimental condition also, Ime1 expression started in YPA, followed by its nuclear localization around 30 mins in SPM with no nuclear localization either in YPA or hardly any in SPM at 10 mins (Fig. S1A, B). FACS analysis showed that the DNA synthesis started before the cells were at least 90 mins in SPM (Fig. S1C), which is consistent with the earlier studies reporting that the initiation of premeiotic S phase in SK1 yeast strain happens around 60 mins in SPM (Borner et al., 2023, Cha et al., 2000). Hence, 1 hr in SPM was taken as the timepoint to harvest meiotic pre-S phase cells for our assays. For metaphase I cells, the sample was harvested at 6 hrs in SPM, and cells with SPB-SPB distance more than 1 but less than 3 μm were analyzed as described before (Tsuchiya et al., 2014, Yu and Koshland, 2005). The mitotic pre-S phase was determined based on Cln2 expression (yeast G1/S-CDK) as its expression marks the beginning of mitosis (Hadwiger et al., 1989) (Fig. S1D).

**Figure 1.**
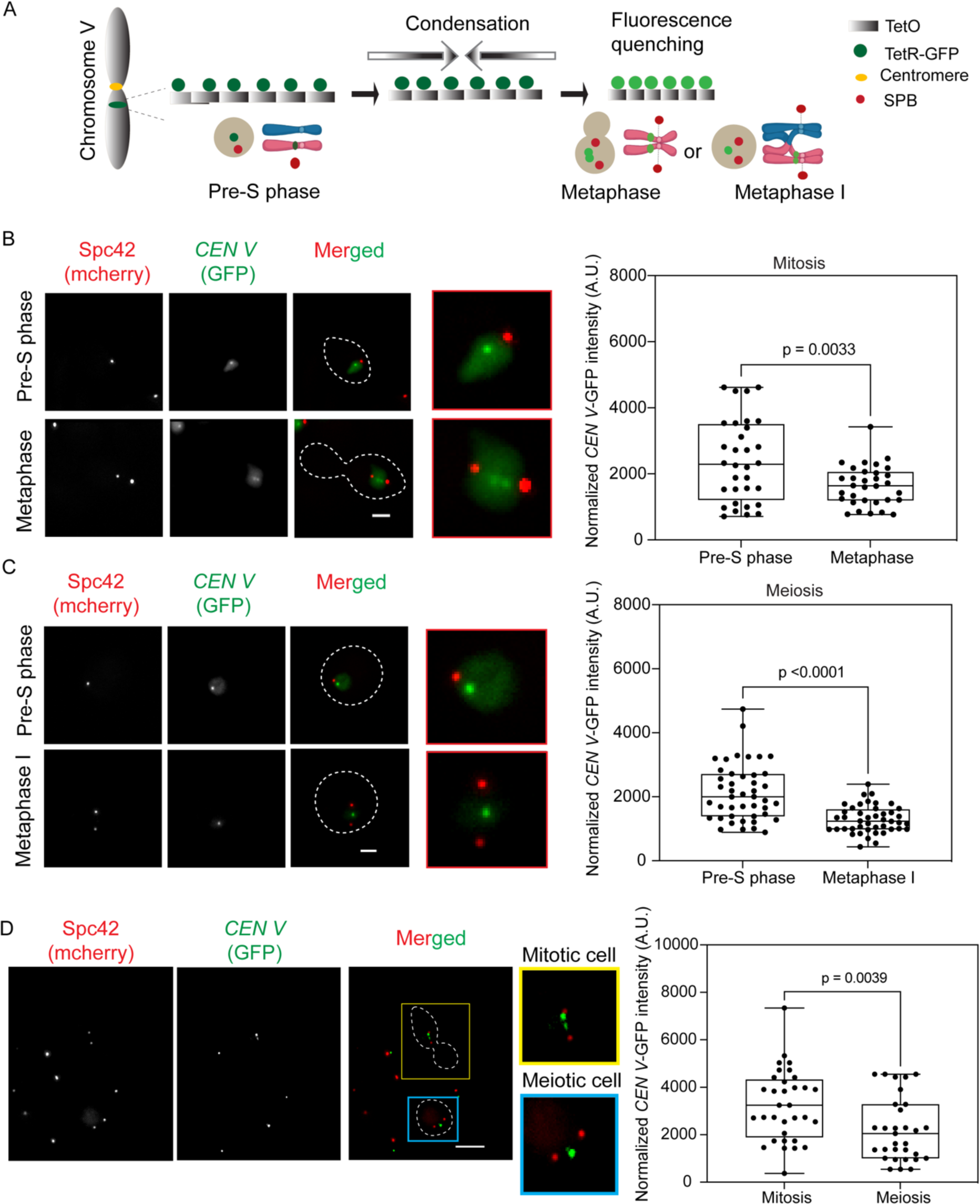
Chromosomes are more condensed in meiosis than in mitosis. (A) Schematic diagrams showing the strategy adopted for in vivo measurement of chromosome condensation. One copy of chromosome V was tagged using [TetO]_224_-[TetR-GFP] system near the centromere V. As the [TetO]_224_ repeats come closer due to axial condensation in the metaphase stage, the GFP signal intensity decreases because of fluorescence quenching. The fluorescence intensity of *CEN V*-GFP measured in (B) mitotic and (C) meiotic cells (see materials methods for detail) from the indicated cell cycle stages in the wild type (SGY9012) are graphically represented on the right; the corresponding representative images are shown on the left. N ≥ 30-45 from three independent experiments, scale bar = 2 µm. (D) The cells used in (B) and (C) were mixed and fluorescence intensity was measured from mitotic (budded), and meiotic (unbudded) cells present in the same field of view as represented on the left; the fluorescence intensities are presented graphically on right. N ≥ 30 from two independent experiments, scale bar = 5 µm. The intensity values were normalized with background intensities. For statistical significance, *p* values were estimated by the two-tailed student’s t-test for the mean.

The *CEN V*-GFP signal intensity normalized with the background intensity was measured from four different types of samples – mitotic interphase, metaphase, meiotic interphase, and metaphase I cells as described in materials and methods. As expected, both in mitosis and meiosis, the *CEN V*-GFP fluorescence intensity decreases in the metaphase and metaphase I cells, respectively, than in the corresponding interphase cells reconfirming that condensation occurs as the cells proceed into the M phase of the cell cycle. (Lavoie et al., 2004; Vas et al., 2007). Interestingly, while in mitosis, the extent of decrease was 31.2% (2389 vs. 1643 A.U., Fig. 1B), the same was 39.6% (2136 vs. 1289 A.U., Fig. 1C) in meiosis. Notably, the signal intensity was observed ∼1.3-fold lesser in meiotic metaphase samples as compared to mitotic metaphase I (1289 vs. 1643 A.U.). Such a decrease in signal intensity in meiosis over mitosis argues for an increased level of chromosome condensation in meiosis than in mitosis. Since *CEN V*-GFP fluorescence intensity was measured from different mitotic and meiotic cells, an error in measuring the intensity across the samples may lead to the observed difference in fluorescence intensity between mitosis and meiosis. To rule out this possibility we mixed the mitotic and meiotic cells and measured the fluorescence intensity from the metaphase or metaphase I cells residing in the same field of view. The metaphase cells with two SPBs and a bud were distinguishable from the metaphase I cells harboring 2 SPBs but no bud. Consistent with the previous observation, we found ∼1.5-fold decreases in signal intensity in meiotic metaphase I cells compared to mitotic metaphase cells (3252 vs. 2198 A.U., Fig. 1D), indicating chromosomes are indeed more condensed in meiosis than in mitosis.

### The non-essential kinetochore protein Ctf19 influences chromosome condensation in meiosis but not in mitosis

Earlier we and others have reported that the mitotically non-essential kinetochore proteins have essential roles in meiosis (Mehta et al., 2014, Agarwal et al., 2015, Borek et al., 2021). In support of this, we observed that such a protein, Ctf19, promotes rDNA condensation in meiosis but not in mitosis (Mehta et al., 2014). Since we revealed that the extent of chromosome condensation, at least at the pericentromeres, is more in meiosis than in mitosis, we wished to investigate the contribution of Ctf19 to this difference. We measured the *CEN V*-GFP signal intensity in the diploid *ctf19Δ* mutant cells from the pre-S phase and metaphase/metaphase I in both mitosis and meiosis. Any meiosis-specific function of Ctf19 on condensation is expected to ameliorate the difference in condensation (or *CEN V-*GFP signal intensity) between the pre-S phase and M phase in meiosis but not in mitosis. We observed a significant (44.2%) decrease in signal intensity in metaphase cells compared to the pre-S phase cells in the *ctf19Δ* mutant (3195 vs 1789 A.U., Fig. 2A) similar to the wild type (Fig. 1B) suggesting that condensation is not compromised in the mutant compared to the wild type in mitosis. In contrast, in meiosis, unlike the wild type, which showed a 39.6% decrease in the intensity in meiosis (Fig. 1C), only around 6.5% decrease (4182 vs 3913 A.U., Fig. 2B) was found in the mutant, which was not statistically significant indicating that a meiosis-specific chromosome condensation defect occurs in absence of Ctf19. Notably, although the interphase intensity was observed higher in *ctf19Δ* than in the wild type irrespective of the type of the cell cycle, the mutant achieves a level of chromatin condensation similar to the wild type during mitotic metaphase, but not during meiotic metaphase I (Fig 2C). This suggests that while the *ctf19Δ* mutation might also have subtle condensation defect in mitosis, its impact is more substantial during meiosis.

**Figure 2.**
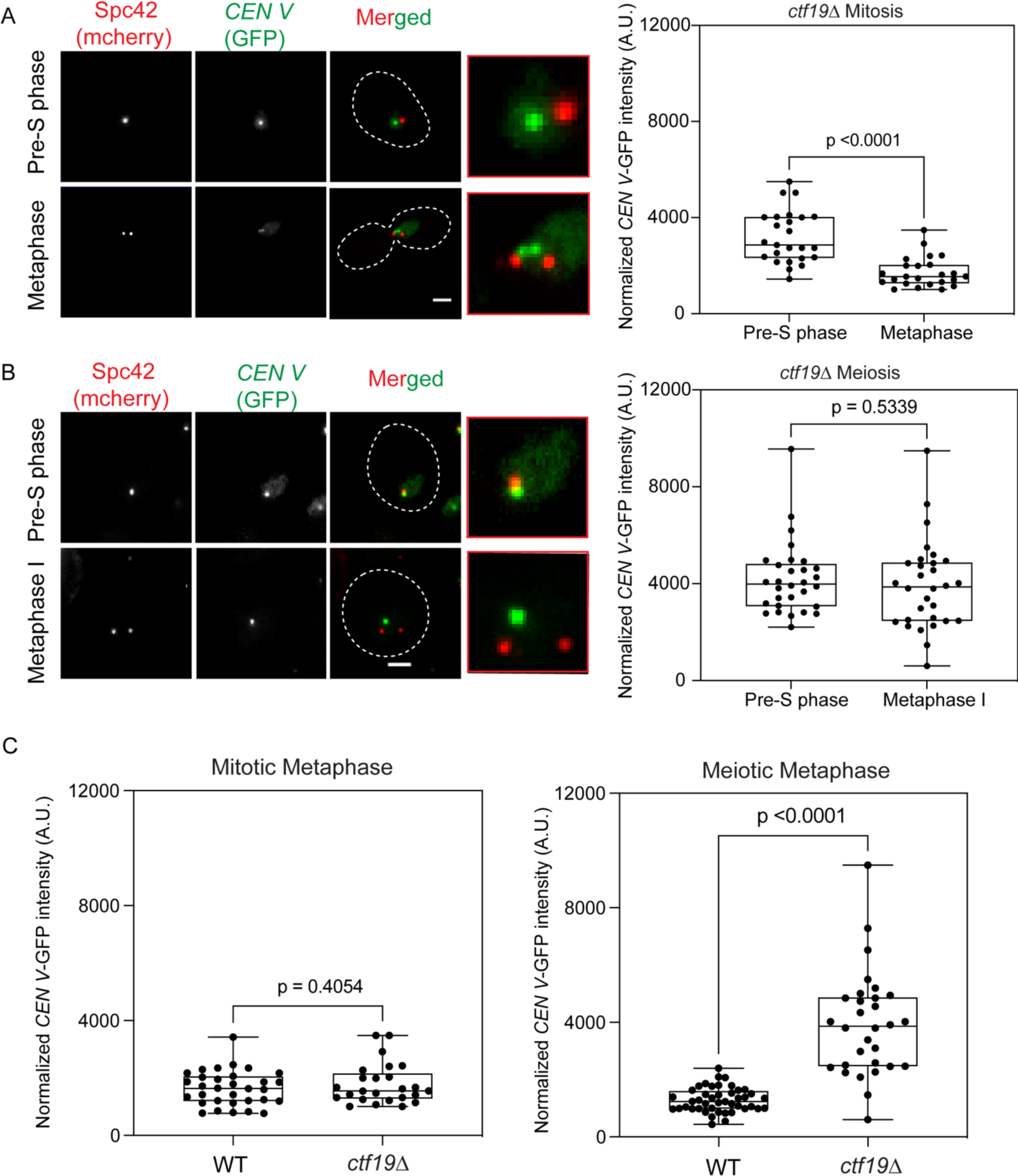
Ctf19 is involved in chromosome condensation in meiosis but not in mitosis. In vivo, chromosome condensation in *ctf19*Δ cells (SGY9019) was measured as in Fig. 1 at the indicated stages in mitosis (A) and in meiosis (B). (C) The fluorescence intensity of *CEN V*-GFP measured for both the wild type (SGY9012) and *ctf19Δ* (SGY9019) cells in the mitotic metaphase stage (left) and meiotic metaphase stage (right) are graphically represented. N = 25-30 from two independent experiments. Error bars represent the standard deviation from the mean values obtained from two-three independent experiments. For statistical significance, *p* values were estimated by the two-tailed student’s t-test for the mean.

To examine if the observed condensation defect in the *ctf19Δ* mutant in meiosis is specific to the Ctf19c proteins of the kinetochore, we monitored the *CEN V*-GFP signal in the pre-S phase and metaphase I stage in the cells lacking Slk19, another non-essential protein residing at the kinetochore but not as a component of the Ctf19c (Zeng et al., 1999). We could not observe any chromosome condensation defect in the *slk19Δ* cells as the GFP signal intensity decreased in metaphase I compared to the interphase stage (∼45%, 592 vs 321 A.U., Fig. S2A). Since we used identical conditions during mitosis and meiosis, the possibility of other factors viz., accessibility of TetR-GFP to the operators or a lower concentration of TetR-GFP responsible for reducing the fluorescence intensity, can be void. Moreover, the control mutant *slk19Δ* behaved as expected.

To further establish that the observed condensation defect in the *ctf19Δ* is not due to an artifact of the assay employed above, we used an alternative in-vivo assay to further validate the observed condensation defect. We evaluated the condensation status in meiosis and mitosis for both the wild type and *ctf19Δ* by using Second Harmonic Generation (SHG) microscopy which relies on generating a second harmonic by the chromatin fibers upon excitation by two-photon microscopy (Yamin et al., 2020, 2022). In this assay, the chromosomes were stained with DAPI and a higher ratio of integrated density to area obtained through the SHG technique corresponds to higher condensation. As expected, we indeed observed an increased condensation in both wild type and *ctf19Δ* in mitotic metaphase as compared to pre-S phase (Fig. 3 A, B). However, when the meiotic cells of the same strains were analyzed, the wild type cells showed significant increase in the ratio of integrated density to area at metaphase I as compared to pre-S phase, but the *ctf19Δ* cells did not show any significant increase in the ratio in meiotic metaphase I (Fig. 3 C, D). These findings further confirm that the *ctf19Δ* cells are indeed compromised in meiotic condensation with no significant effect on mitotic condensation.

**Figure 3.**
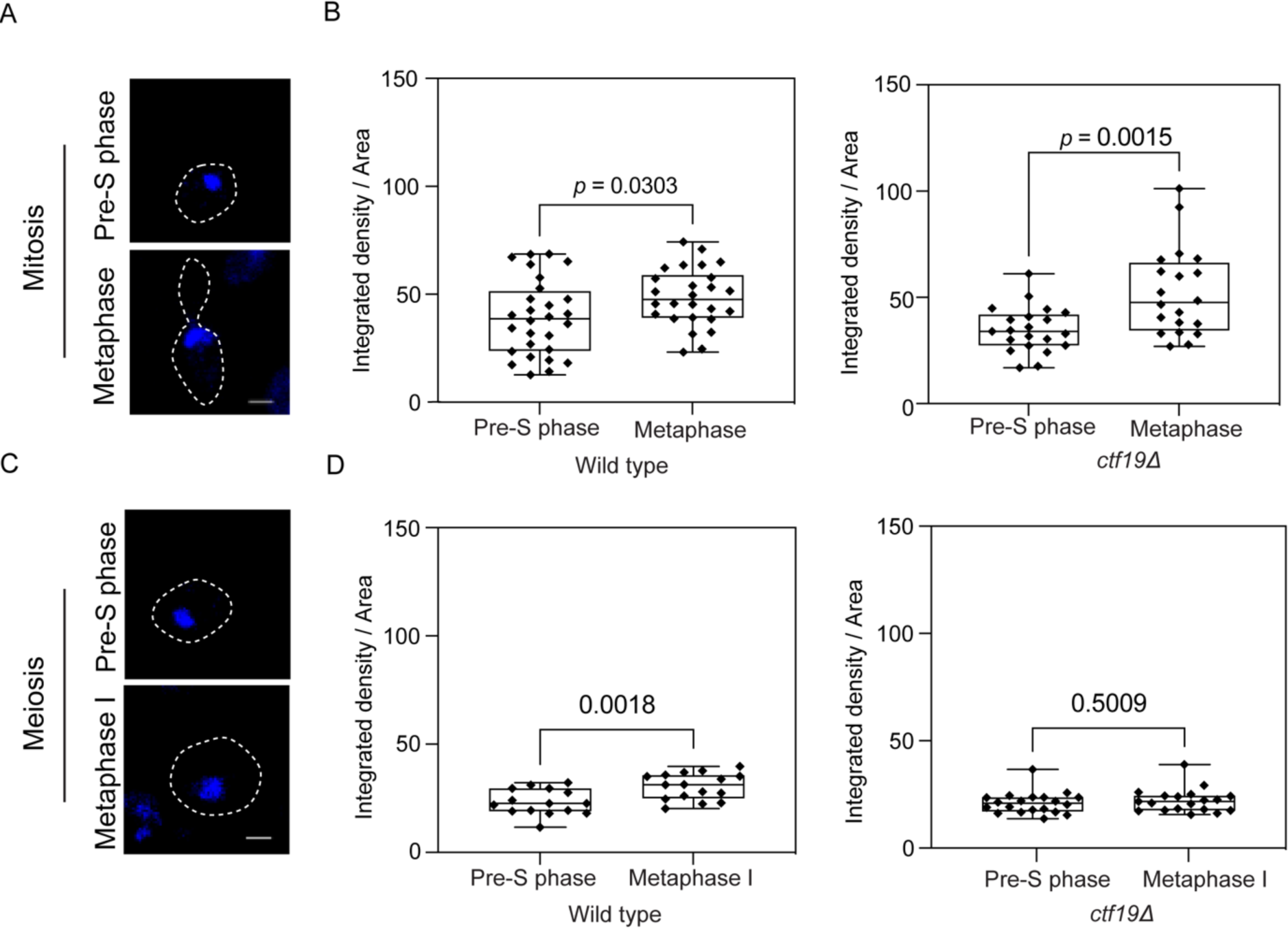
The loss of Ctf19 leads to meiotic specific chromosome condensation defect. (A) Analysis of condensation of the chromosomes stained with DAPI in wild type (SGY9012) and *ctf19Δ* (SGY9019) mitotic cells by Second Harmonic Generation (SHG) microscopy. Cells in the mitotic pre-S phase or metaphase were selected based on bud morphology and DAPI position within the cell. (B) Box plot showing the integrated density/area from pre-S phase cells and M phase cells after analysis using Image J for the indicated strains. (C) Similar analysis as in A for the meiotic cells. Cells in meiotic pre-S phase or metaphase I were collected at different time points following their release into SPM as mentioned in Fig S1 (see materials and methods). (D) Similar graphs as in B for the meiotic cells. Representative images show the areas taken for analysis after two-photon excitation in SHG microscopy.

Since in budding yeast the rDNA condensation can be monitored in a better way than the rest of the chromosomes, for further validation of the condensation defect in the *ctf19* mutant, we compared the extent of rDNA condensation using Net1-GFP as the marker as described before (Torres-Rosell et al., 2004, Machín et al, 2005, Mehta et al., 2014) among the wild type, *ctf19Δ*, and *slk19Δ* strains (Fig. S2B). The Net1-GFP signal pattern was visualized in these strains when the cells were arrested at the metaphase I stage (*pCLB2-CDC20*). While around 85% of the wild-type cells showed the ‘tight knit’ and ‘crescent’ signal which represent a condensed state of rDNA (Mehta et al., 2014; Kerr et al., 2011), *ctf19Δ* cells showed a reduction (50%) in the frequency and a corresponding increase (50%) in the occurrence of the ‘puff’ signal which corroborates a decondensed state. On the other hand, *slk19Δ* cells showed similar results to the wild type. In mitosis, however, there was no difference in condensation amongst the wild type, *ctf19Δ*, and *slk19Δ* cells (Fig. S2C) where the ‘line’, ‘loop’, and ‘puff’ patterns designate condensed, intermediate, and decondensed forms of the rDNA locus, respectively (D’Ambrosio et al., 2008; Dauban et al., 2019). Altogether, from these results, we infer that Ctf19 promotes chromosome condensation in meiosis but not in mitosis, and this function appears to be specific for the Ctf19 protein or perhaps for all the non-essential proteins of the Ctf19c of the kinetochore.

### Loss of Ctf19 affects axial condensation of the chromosomal arm

The loss of pericentric and rDNA condensation in the *ctf19*Δ mutant prompted us to examine the mutant for linear (axial) condensation of the chromosomal arms which relies on condensin (Kruitwagen et al., 2015). To investigate this, we marked chromosomes at two different loci 400 kb away from each other, one was near the *CEN V* (1.4 kb away), as before, and the other one was 21 kb away from *TEL V* using [TetO]_224_ –[TetR-GFP] and [TetO]_448_–[TetR-GFP] systems, respectively. Since the number of operators used to visualize *TEL V-*GFP was double that of *CEN V*-GFP, the two loci could be visually distinguishable by the brightness of the GFP dots where *TEL V*-GFP appeared much brighter than the *CEN V*-GFP (Fig. 4A, B). In the wild type, we observed a 37% decrease in the mean distance in the metaphase I stage (0.568 µm) compared to the pre-S phase stage (0.83 µm) owing to the condensation (Fig. 4C, left panel). However, in the *ctf19Δ* mutant, there was no significant decrease in the mean distances between the two stages (1.00 vs. 1.167 µm), suggesting a defect in the condensation (Fig. 4C, right panel). While looking at the axial condensation in mitosis, there was a statistically significant decrease in the distance in metaphase compared to the pre-S phase both in the wild type and the mutant (21% and 24 %, respectively) (Fig. S3 A, B). Notably, since the mean distance in the mutant (1.16 µm) was found higher than the wild type (0.83 µm) in the meiotic pre-S phase that precedes condensin function, it is possible that at this phase, the compaction of the chromatin is compromised in the mutant due to relaxed nucleosome-nucleosome interaction or probably due to subtle defects acquired during preceding mitotic cycle. Altogether, these results suggest that the contribution of Ctf19 in promoting chromatin condensation in meiosis is not only restricted to the pericentromeric regions but also extends axially along the chromosome arms at the distal loci.

**Figure 4.**
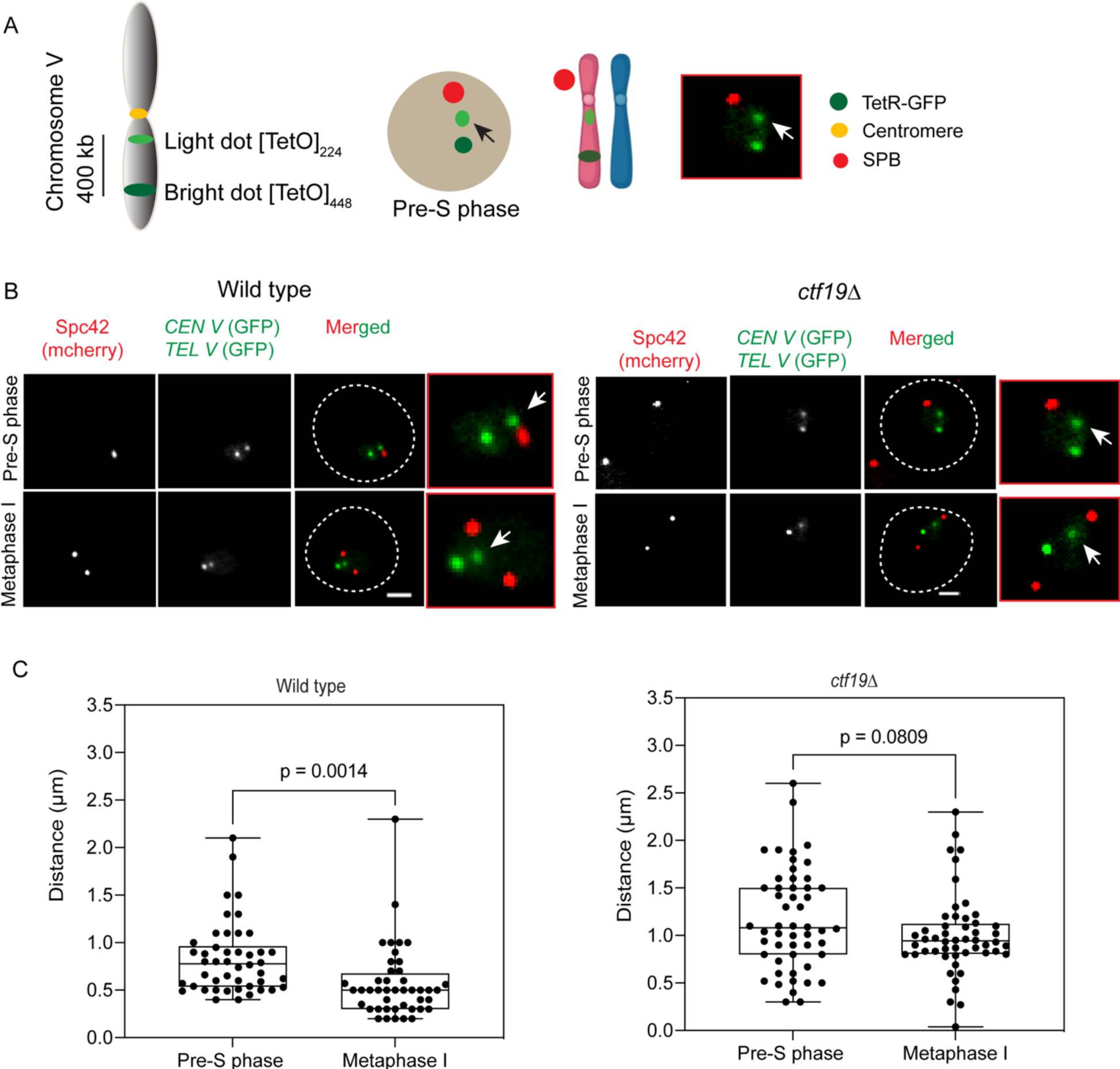
Ctf19 contributes to chromosome condensation along the chromosomal arm. (**A**) One copy of chromosome V was marked with GFP at two different chromosomal loci ∼400 kb away from each other, each proximal to the centromere or telomere was marked with [TetO]_224_–[TetR-GFP] as *CEN V*-GFP or with [TetO]448 –[TetR-GFP] as *TEL V*-GFP, respectively. The *TEL V-*GFP dot appeared brighter than the *CEN V*-GFP dot (arrow). (B) The representative images are shown for the indicated strains (wild type: SGY9092, *ctf19Δ*: SGY9124) and the cell cycle stages. Scale bar = 2 µm. (C) The 3D distances between the *CEN V*-GFP and *TEL V*-GFP dots measured as described in the materials and methods are shown for the wild type (left) and *ctf19*Δ (right) strains used in B harvested from the indicated cell cycle stages. N = 45-50 from two independent experiments. The statistical significance *p* value was estimated by the two-tailed student’s t-test for the mean.

### Ctf19 is required for maintenance of chromatin-condensin association in meiosis

Earlier, we reported that in the absence of Ctf19, the chromatin-condensin association is perturbed in meiosis (Mehta et al., 2014), which supports our findings from the condensation assays described here. We wished to re-verify our earlier observation with further investigation to find whether the effect on chromatin-condensin association is specific to Ctf19c proteins and whether the effect is on the establishment and/or maintenance of the association. To address these, we first compared the localization of the condensin protein Brn1 on the chromatin between the wild type and *ctf19Δ* mutant along with the *slk19Δ* mutant, which, as a control, showed no effect on condensation (Fig. S2A and S2B). Brn1 fused to 6XHA at the C-terminal (Verzijlbergen et al., 2014) was detected on the chromatin in the meiotic cells arrested at metaphase I by Cdc20 depletion. While a similar number (∼ 13 to 15) of bright Brn1-6HA foci were visible in the wild type and *slk19Δ* mutant, the number of foci was much lower (∼7), and the foci were less bright in the *ctf19Δ* mutant suggesting a defect in chromatin-condensin association in the latter (Fig. S4 A). On the other hand, as expected from the condensation assay results, no difference in Brn1-6HA localization was observed between the wild type and *ctf19*Δ spreads prepared from the cells arrested at mitotic metaphase by Cdc20 destruction as mentioned in materials and methods (Fig. S4 B, C). To test if the observed condensation defect is specific to Ctf19 or also occurs in other non-essential proteins of Ctf19c, we analyzed the association of Brn1-6HA with chromatin in cells lacking another protein of Ctf19c, Ctf3. We found that in *ctf3Δ* cells also, the number of Brn1-6HA foci was lesser (∼4) than the wild type (∼9) (Fig. S5). The lesser number of foci here in the wild type compared to the previous chromatin spread of Brn1-6HA in the wild type (Fig. S4) can be because of technical difference in antibody staining, as these experiments were performed at different times To further validate our chromatin spread data from meiosis, we performed a ChIP assay to quantify the association of Brn1 with different regions of chromosome – centromere, pericentromere, and arm (Fig. 5A). The cells were harvested after 8 and 10 hrs into SPM to obtain metaphase I arrested wild type and *ctf19Δ* mutant, respectively by Cdc20 depletion. The time points were chosen based on our previous observations where the *ctf19Δ* cells showed a delay in the meiotic cell cycle and were arrested maximally at metaphase I at 10 hrs into SPM (Mehta et al., 2014). We observed a significant reduction in the level of Brn1 at all the regions in the mutant compared to the wild type; however, the reduction was more at the centromeric region compared to other regions (Fig. 5B). These results, re-confirming our earlier chromatin spread data (Mehta et al., 2014), further indicate that the meiotic defect in chromatin-condensin association is specific to the lack of Ctf19c proteins.

**Figure 5.**
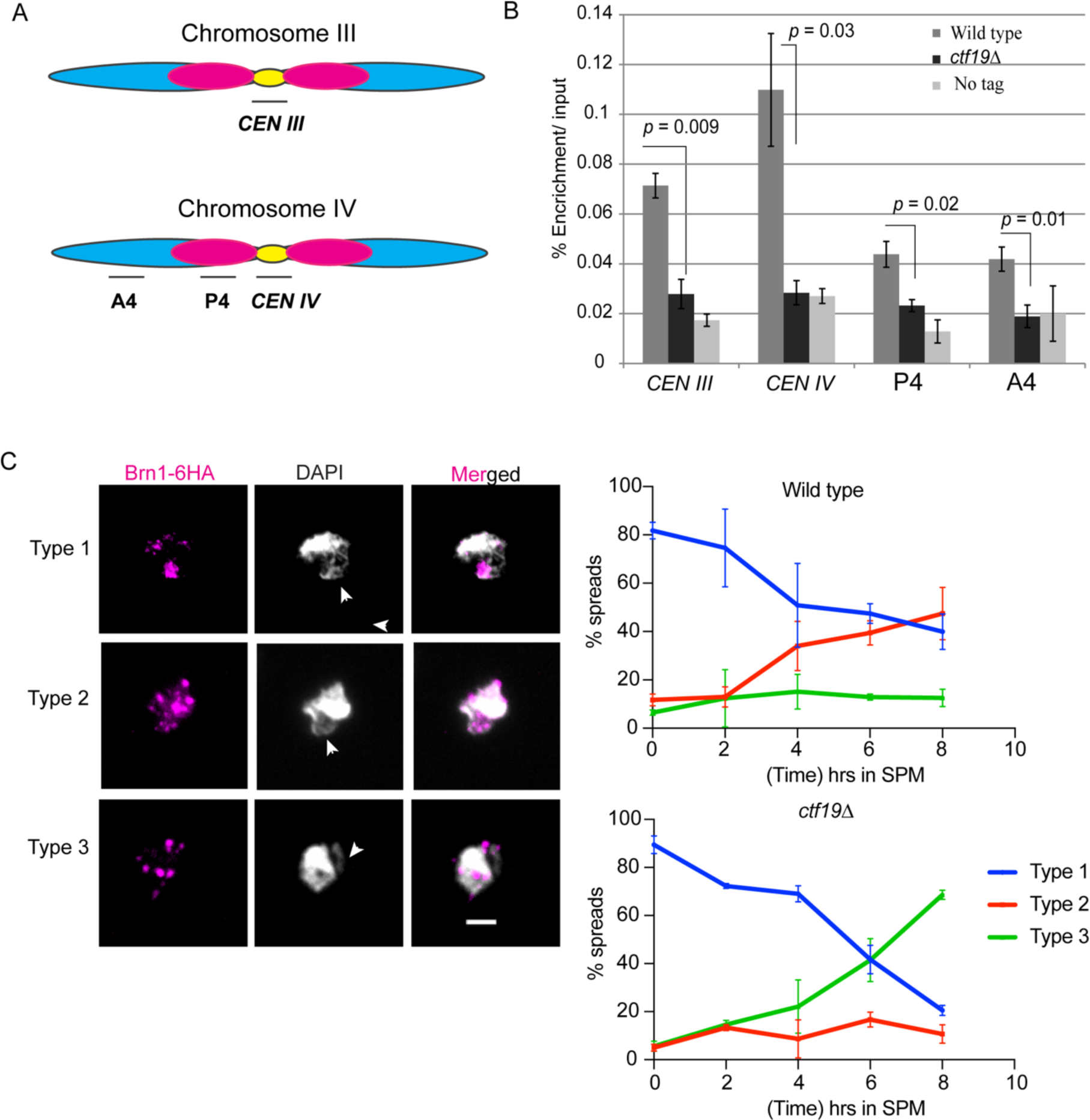
The loss of Ctf19 causes reduced association of condensin with chromatin by impairing the establishment of the association in meiosis. (A) Schematic diagram showing the locations of PCR amplicons (horizontal bars) on chromosomes III and IV used for the qPCR analysis of the ChIP samples. The yellow oval shapes represent the centromeres (*CEN III* and *CEN IV*), pink and blue regions indicate the pericentromeric (P) and arm (A) regions, respectively. (B) ChIP assay using anti-HA antibodies (3F10) for quantifying the association of Brn1-6HA with the indicated chromosomal loci in the wild type (SGY283) and *ctf19Δ* (SGY284) cells arrested at metaphase I. The graph represents % enrichment per input obtained by qPCR analysis. Error bars represent the standard deviation from the mean values obtained from three independent experiments. The statistical significance *p* value was estimated by the two-tailed student’s t-test for the mean. (C) The loss of Ctf19 impairs the establishment of the association of condensin with chromatin in meiosis. The left panel shows the representative images of the chromatin spreads showing different types of association (described in the text) of the condensin subunit (Brn1-6HA) with chromatin at different time points in SPM in strains used in B harboring homozygous *BRN1-6HA* allele. Types 1 – 2 are categorized as localization of Brn1-6HA first at rDNA (type 1) and then gradually spreading to rest of the chromatin (type 2). Type 3 implies a gross reduction in chromatin-condensin association. Anti-HA antibodies (3F10) were used to detect Brn1-6HA. The right panel represents the percentage of different types of spreads at the indicated time points. with respect to the Brn1-6HA signal. Arrowhead shows rDNA region. N >100 from two independent experiments, scale bar = 2 µm.

Next, we wished to address whether Ctf19 is required for the establishment or maintenance of chromatin-condensin association. It has been previously shown that during early meiosis, condensin first associates with rDNA as a cluster and then at later time points associates with the rest of the chromatin, leaving a minuscule amount at the rDNA (Markowitz et al., 2020; Li et al., 2014). To compare the dynamics of chromatin-condensin association between wild type and *ctf19Δ* we monitored the association of Brn1-6HA with chromatin using chromatin spreads at different time points in the cells released into synchronized meiosis (Fig. 5C). Types 1 – 2 are categorized as localization of Brn1-6HA first at rDNA (type 1) and then gradually spreading to rest of the chromatin (type 2). Type 3 implies a gross reduction in chromatin-condensin association. While the initial rDNA localization of Brn1-6HA (0 hr, type 1) in the mutant was not found lesser than the wild type, its gross chromatin localization (type 2) in the mutant was merely below 10% of the spreads compared to the wild type (40%) at 4 hrs. The spreads with type 2 localization of Brn1 did not increase in the mutant even at the later time points (∼10% at 8 hrs), while the spreads from wild type cells showed about ∼50% type 2 localization at the same time points. This implies that the establishment of condensin association with the gross chromatin, but not with the rDNA, is perturbed in the absence of Ctf19. Consequently, at 8 hrs, while the mutant showed a major population of spreads (70%) with reduced condensin on the chromatin (type 3), only a minor population (<10%) of wild type spreads demonstrated this phenotype. The phenotype of reduced chromatin association of Brn1 (type 3) in the mutant starts to appear as early as 4 hrs and keeps on increasing as the cells undergo the meiotic cell cycle (Fig. 5C). Moreover, the type 1 population also decreased to <10% in the mutant, while about 50% of the wild type cells retained the type 1 population at 8 hrs; these observations may be interpreted as a defect in the maintenance of the rDNA association of condensin in the mutant.

### The condensation defect in *ctf19Δ* may occur partially through Sgo1 mislocalization but independent of DDK- and Ipl1-mediated pathways

Next, we wished to address the mechanism through which Ctf19 may regulate condensation in meiosis. Besides protecting the pericentromeric cohesin in meiosis I, Sgo1 (Shugoshin) has a role in mitotic chromosome condensation (Kruitwagen et al., 2018) through condensin loading at pericentromeric region (Peplowska et al., 2014). It has also been reported that in the absence of Ctf19c proteins, Iml3 and Chl4, the Sgo1 localization becomes compromised at the chromatin both in mitosis (Deng et al., 2018) and meiosis (Kiburz et al., 2005). To investigate if Ctf19, unlike the peripheral Ctf19c components Iml3 and Chl4, differs in recruiting Sgo1 on the chromatin between mitosis and meiosis which may cause the differential effect on condensation, we analyzed the localization of Sgo1-EGFP in mitosis (metaphase) and meiosis (metaphase I) in presence or absence of Ctf19. We observed that the absence of Ctf19 affected Sgo1 localization both in mitosis (Fig. 6A) and meiosis (Fig. 6B), however, the extent of the defect was more in meiosis where 81% of the cells showed Sgo1-EGFP mislocalization (no signal) compared to only 57% in mitosis. Whereas the fraction of cells showing no signal of Sgo1-EGP in the wild type was found similar in mitosis and meiosis (Fig. 6A, B). To further validate our cell biological data, we performed a ChIP assay to quantify the association of Sgo1-9Myc with the centromeres. We observed a significant reduction in the level of Sgo1-9Myc at the centromeres both in mitosis (Fig. 6C) and meiosis (Fig. 6D) in *ctf19Δ* mutant compared to the wild type, while the reduction was non-significant at a negative locus, *TUB2.* However, the reduction was more (∼86% (0.2114 vs 0.03) for *CEN III* and ∼82% (0.2497 vs 0.043) for *CEN IV*) in meiosis than in mitosis (∼ 45% (0.4290 vs 0.2376) for *CEN III* and ∼ 40% (0.3745 vs 0.2254) for *CEN IV*) (Fig. 6C, D; see materials and methods). Based on these observations, we conclude that a higher reduction in Sgo1’s association with the chromatin in meiosis over mitosis in *ctf19Δ* cells may contribute partially, if at all, to the meiosis-specific chromatin-condensin association defect in these cells.

**Figure 6.**
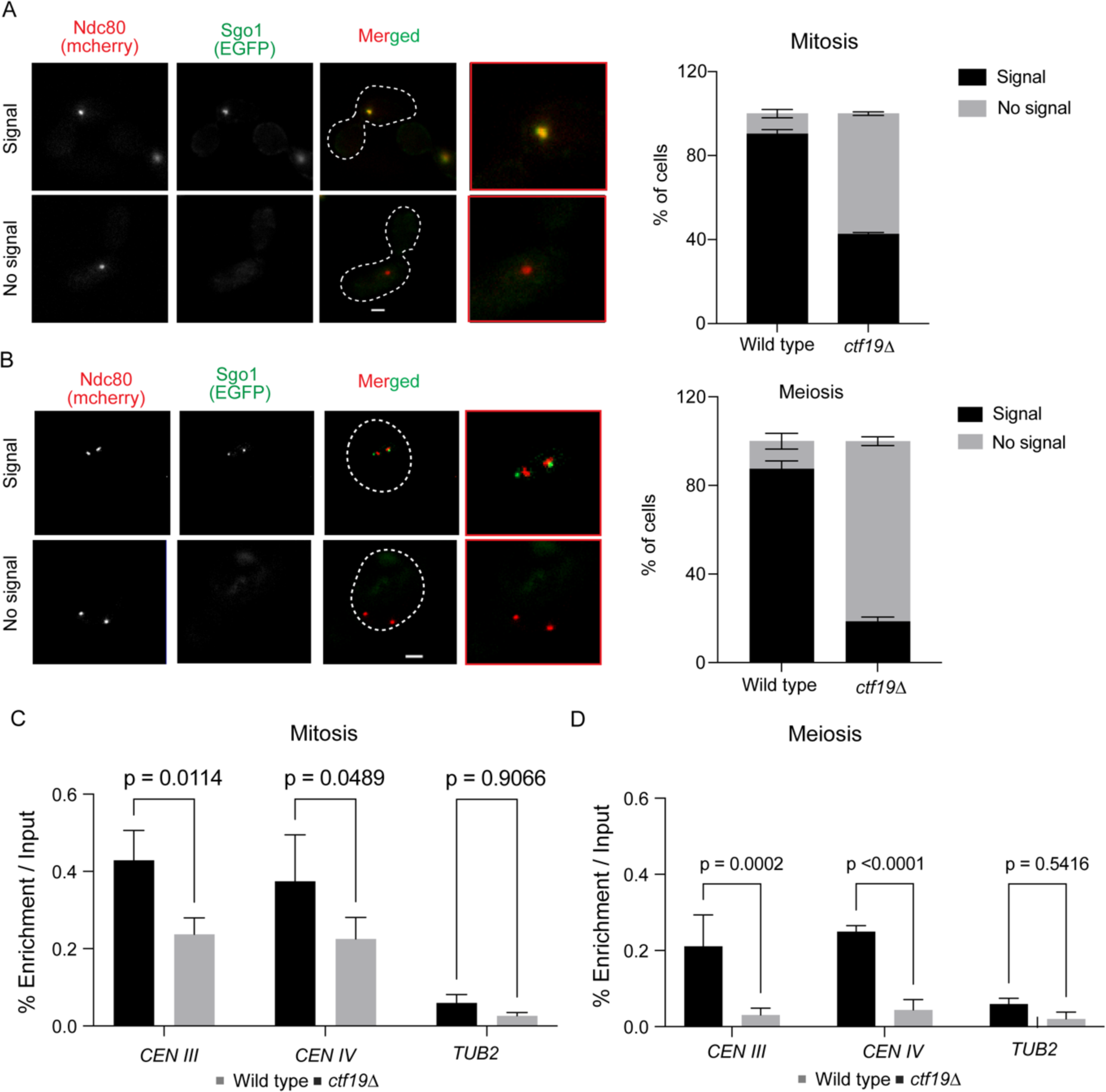
Pericentric localization of Sgo1 is perturbed in the *ctf19Δ* cells compared to the wild type more in meiosis than in mitosis. The localization of Sgo1-GFP with respect to a kinetochore protein, Ndc80-mCherry in the wild type (SGY9189) and *ctf19Δ* (SGY9193) cells arrested at metaphase upon treatment with nocodazole (A) and harvested at metaphase I following their release into SPM for 8 and 10 hrs (B), respectively. The left panel shows the representative images of indicated categories and the right panel depicts the percentage of cells in each category. Error bars represent the standard deviation from the mean values obtained from two independent experiments. N = 90-120, scale bar = 2 μm. ChIP assays using anti-Myc antibodies (ab9106) for quantifying association of Sgo1-9Myc with the indicated chromosomal loci in the wild type (SGY9194) and *ctf19Δ* (SGY9195) cells arrested at metaphase (C) and metaphase I (D), as in (A) and (B). The graph represents % enrichment per input obtained by qPCR analysis. Error bars represent the standard deviation from the mean values obtained from three independent experiments. The statistical significance *p* value was estimated by the two-tailed student’s t-test for the mean.

It has been reported earlier that till metaphase the rDNA condensation in mitosis is dependent on cohesin (Lavoie *et al*., 2002). As the cohesin loading at the pericentromeres is dependent on Ctf19 through recruitment of DDK (Dbf4-dependent kinase) both in mitosis and meiosis (Hinshaw et al., 2015, 2017), therefore, the reduction of cohesin level at the centromeres in *ctf19*Δ mutant is unlikely the cause for observed meiosis-specific condensation defect. However, recently DDK and Cdc5 (budding yeast polo-like kinase) have been reported to have roles in the meiotic prophase-like pathway (cleavage-independent cohesin removal from the chromatin) which has been shown important for chromosome compaction in meiosis probably by making space on chromatin for condensin loading (Challa et al., 2019). Based on this, we hypothesized that perhaps DDK kinase localization at the centromeres is perturbed in the absence of Ctf19, which in turn results in an impaired prophase-like pathway, hence leading to chromosome condensation defect. If our hypothesis is correct, then a DDK mutant unable to be localized at the centromeres should show a condensation defect in meiosis. It has been reported that the fusion of epitopes at the C-terminal of the Dbf4 protein renders it inactive at the centromeres however, its localization at the origins of replications remains unaffected (Hinshaw et al., 2017). We constructed a strain expressing Dbf4-9myc with heterozygous tagging of *CEN V* and *TEL V* with GFP. We confirmed that the *DBF4-9MYC* allele is acting as a mutant, as we observed a reduced frequency of sister chromatid cohesion (Fig. S6A) as reported earlier (Hinshaw et al., 2015). However, we failed to observe any defect either in axial contraction by measuring the distance between *CEN V* and *TEL V* (Fig. S6B) or in condensation of centromeric and telomeric regions measured by fluorescence-based quenching assay (Fig. S6C). Here, in meiosis, although we observed that the distances between the two loci are more than the wild type (Fig 4B), but the extent of axial compaction from pre-S phase to metaphase I was found similar in both the conditions viz., ∼1.4 fold in the wild type (0.8308 vs 0.5686 μm) while ∼1.2 fold in the DDK mutant (1.179 vs 0.9591 μm). Based on these observations we conclude that the absence of DDK at the centromeres in *ctf19Δ* cells is not responsible for the observed condensation defect in these cells in meiosis.

Phosphorylation of histone H3 at serine 10 by Aurora kinase B (Ipl1 in budding yeast) has been known to play a crucial role in chromosome compaction both in yeast (Kruitwagen et al., 2015) and mammals (Mora-Bermúdez et al., 2007). Additionally, an in vitro assay has demonstrated an interaction between Ctf19 and Ipl1, which is essential for stabilizing the chromosomal passenger complex (CPC) at the centromeres (Fischböck-Halwachs et al., 2019). However, it is uncertain whether Ctf19 interacts with Ipl1 in vivo and regulates Ipl1’s centromeric localization and possibly thereby promotes chromosome condensation in meiosis. To address this question, we performed a ChIP assay to quantify the association of Ipl1 at the centromeres in metaphase I arrested wild-type and *ctf19*Δ mutant cells, harvested at 8 and 10 hours into SPM, respectively. However, our analysis did not reveal any significant reduction in the centromeric enrichment of Ipl1 in the *ctf19Δ* mutant compared to the wild type (Fig. S7). Based on these findings, we conclude that Ctf19 does not regulate Ipl1 localization at the centromeres, at least during meiosis, and hence Ctf19’s involvement in chromosome condensation is independent of the Ipl1-mediated pathway.

### Polo-like kinase (Cdc5) level increases in the cells in the absence of Ctf19 in meiosis

Having shown that DDK and Ipl1 kinases are not involved, we wished to identify other factor(s) that may promote meiotic condensation via Ctf19. A kinase switch model for chromosome condensation says that different kinases take turns to regulate the condensation at different stages of the cell cycle (Bazile et al., 2010). For instance, casein kinase II, cyclin-dependent kinase, and polo-like kinase regulate condensation at the interphase, metaphase, and anaphase, respectively. Recently, in budding yeast, the polo-like kinase Cdc5 has been shown to be important for both condensation establishment and maintenance in mitosis (Lamothe et al., 2020). Furthermore, earlier, we reported that in meiosis, Ctf19c protein mutants show SPB misduplication and supernumerary SPBs (Agarwal et al., 2015), a phenotype also noticed in meiotic cells overexpressing Cdc5 (Shirk et al., 2011). Therefore, we wished to examine if there is any meiosis-specific misregulation of Cdc5 in the *ctf19Δ* cells, which might cause the condensation defect. Surprisingly, we observed a higher level of Cdc5 in the *ctf19*Δ cells in meiosis but not in mitosis when the cells were arrested at metaphase I and metaphase, respectively (Fig. 7A and Fig. S8A). To investigate if the high level is due to increased stability of Cdc5 and/or high expression of the corresponding gene, we first detected the protein level of Cdc5-6HA over time in meiosis after blocking the protein synthesis using cycloheximide (CHX) which was added at 8 hrs and 10 hrs timepoints when the wild type and mutant cells were into SPM medium, respectively. We found that Cdc5 is more stable in the mutant compared to the wild type (Fig. 7B). The difference between the slopes of the lines was significant as calculated using GraphPad Prism 8.0.1 (p = 0.009). However, we failed to observe any increased stability of Cdc5 in the mutant in mitosis (Fig. S8B). To examine if the higher level of Cdc5 in the mutant is due to transcriptional alteration of *CDC5* gene, we measured its transcript level in the wild type and *ctf19Δ* cells used in Fig. 7A. We did not observe any increased expression of the *CDC5* gene in the mutant (Fig. S8C). From these results, we conclude that perhaps increased stability and a consequent higher level of Cdc5 in *ctf19*Δ cells in meiosis cause the condensation defect.

**Figure 7.**
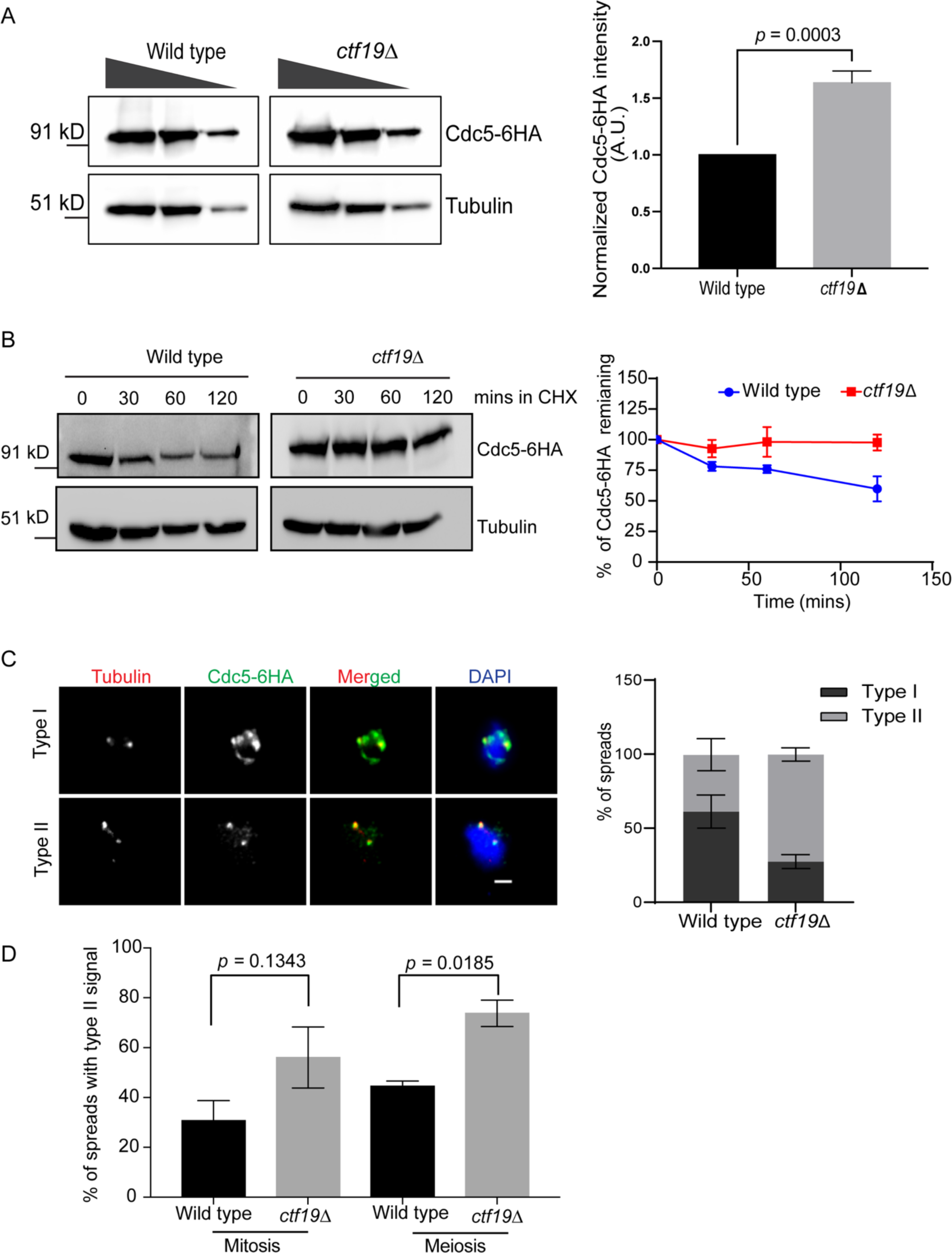
Increase in the level of Cdc5 and decrease in its chromatin association in *ctf19Δ* mutant in meiosis. (A) The wild type (SGY9152) and *ctf19Δ* (SGY9148) cells harboring *CDC5-6HA pCLB2-CDC20* were arrested at the metaphase I following 8 and 10 hrs of meiotic induction, respectively. Left, the proteins extracted from the cells were run on a gel in dilutions and were detected by western blotting using anti-HA (3F10) and anti-tubulin (YOL1/34) antibodies. Right, the intensity of the bands was calculated using ImageJ software and normalized to the tubulin band intensity used as loading control. *p* value was estimated by the two-tailed student’s t-test for the mean (B) Cdc5-6HA stability in wild type (SGY9139) and *ctf19*Δ (SGY9140) cells. Following release into synchronized meiosis for 8 and 10 hrs for the wild type and the mutant, respectively, the cells were treated with CHX (t = 0). The cells were harvested at indicated time points. Cdc5-6HA and tubulin levels were analyzed by western blotting (*left*) as in (A) and quantified (*right*) at indicated time points after CHX treatment. Error bars represent the standard deviation from the mean values obtained from three independent experiments. (C) Left, the representative images of the chromatin spreads show the association of Cdc5-6HA with chromatin in metaphase I arrested wild type and *ctf19Δ* strains as used in A harboring homozygous *CDC5-6HA* alleles. Right, quantitative analysis of Cdc5-6HA signal (*right*) on the chromatin spreads as type I (bright signal) and type II (low signal) categories. The metaphase I stage of the spreads was determined by SPB localization revealed by tubulin immunostaining. (D) The percentage of chromatin spreads with type II signal, as mentioned in (C), from the cells arrested in metaphase (mitosis) by Cdc20 degradation (materials and methods) or metaphase I (meiosis). Error bars represent the standard deviation from the mean values obtained from two independent experiments (N = 83, scale bar = 2 µm).

Despite the high level of Cdc5, its association with chromatin was reduced in *ctf19*Δ mutant compared to the wild type as observed by chromatin spread in meiosis (Fig. 7C). We observed a significant increase in the population of spreads with reduced Cdc5 signal (type II) in the *ctf19Δ* strain compared to the wild type during meiosis. However, this difference was not significant during mitosis (Fig. 7D). The reduction in Cdc5-chromatin association can be possible because of reduced cohesin association with chromatin in the absence of Ctf19 (Mehta et al., 2014, Hinshaw et al., 2017). In the absence of cohesin, the chromatin localization of Cdc5 is compromised and its activity becomes somehow misregulated causing hyper-phosphorylation of condensin subunits leading to condensation defect (Lamothe et al., 2020). Therefore, we hypothesized that the increased level of chromatin unbound Cdc5 in *ctf19*Δ meiotic cells may misregulate the condensin phosphorylation status resulting in a reduced association of condensin with the chromatin with consequent condensation defect (Fig. 5). To test this hypothesis, we monitored the phosphorylation status of the condensin protein Ycg1, as it is an essential subunit for condensin complex recruitment to chromatin (Piazza et al., 2014) and is known to be phosphorylated at multiple sites (St-Pierre et al., 2009). We monitored Ycg1-6HA expression pattern over different time points in cells released synchronously into meiosis till metaphase I arrest (Fig. 8A). At later time points (8 hrs and 10 hrs) in SPM, which corresponded to the metaphase I arrested stage, two bands of higher (B1) and lower (B2) mobilities were visible while at early time points (0 and 6 hrs) only one band was visible both in wild type and mutant indicating post-translational modification of Ycg1-6HA as reported earlier (St-Pierre et al., 2009). Since the B1 bands at 6 hrs are of higher intensity compared to the bands at 0 hr, we speculated that 0 and 6 hrs bands harbor unmodified and (unmodified + modified) forms of Ycg1-6HA, respectively. However, it was interesting to note that in the mutant the top band B2 was reproducibly of higher intensity compared to the wild type. We speculated that the B2 band depicts a higher-level modification of Ycg1-6HA and was found more in the mutant compared to the wild type. Treatment of the protein extract from metaphase I arrested cells with lambda phosphatase showed that the modification is phosphorylation (Fig. 8B). A better resolution of the protein bands using Phos-tag gel revealed that B1 band indeed contained unphosphorylated and phosphorylated forms of Ycg1-6HA whereas B2 band harbored hyper-phosphorylated Ycg1-6HA (Fig. 8B). These results suggest that hyper-phosphorylation of condensin protein in the mutant possibly occurs due to excess Cdc5 level in the mutant as observed earlier.

**Figure 8.**
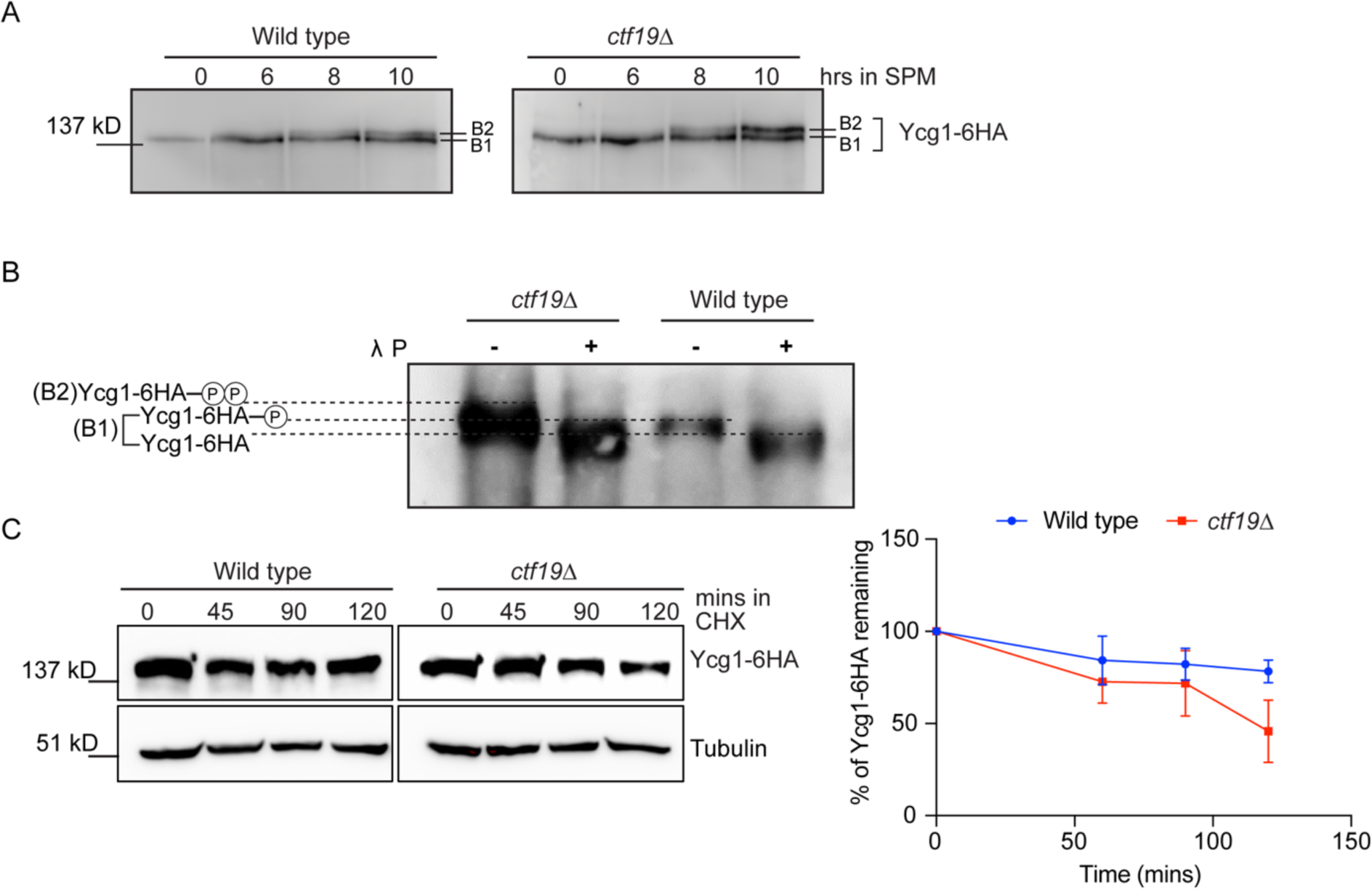
Evidence of hyper-phosphorylation and reduced stability of Ycg1 in *ctf19* mutant in meiosis. (A) The wild type (SGY9232) and *ctf19Δ* (SGY9233) cells harboring *YCG1-6HA pCLB2-CDC20* were released into synchronized meiosis and were harvested at the indicated time points; proteins extracted by the TCA method (materials and methods) from the cells were resolved by running 6% SDS-PAGE gel and were detected by western blotting using anti-HA antibodies (12CA5). Notably, the B2 band is more prominent than the B1 band in the mutant. (B) The wild type and *ctf19Δ* strains used in A harboring *YCG1-6HA pCLB2-CDC20* were released into SPM and the samples were harvested at 8 hrs and 10 hrs, respectively. Protein extracts treated with (+) or without (-) lambda phosphatase (λ P) were run on 7.5% SDS gel made with Phos-tag acrylamide (Wako) to detect Ycg1-6HA by western blotting using anti-HA antibodies (12CA5). (C) Ycg1-6HA stability in wild type and *ctf19*Δ cells. The cells were treated with CHX (t = 0) following 8 and 10 hrs into meiotic induction for the wild type and *ctf19Δ*, respectively. Left, the protein extracts were resolved on 10% SDS-PAGE gel and the protein level was analyzed by western blotting using anti-HA (12CA5) and anti-tubulin (YOL1/34) antibodies in the cells harvested at indicated time points. Right, quantification of Ycg1-6HA bands normalized with the tubulin bands using ImageJ software at indicated time points after CHX treatment. Error bars represent the standard deviation from the mean values obtained from two independent experiments.

It is possible that the hyper-phosphorylated form of Ycg1 in the mutant is less stable which impedes the chromatin-condensin association in the mutant. Similar effect on the chromatin-condensin association was observed for Brn1 subunit of the condensin complex (Fig. 5). To verify this, we monitored the Ycg1 level in the CHX chase experiment where the protein synthesis was blocked by adding the drug at 8 hrs and 10 hrs in SPM for the wild type and the mutant, respectively; the samples were harvested at the indicated time points (Fig. 8C). We found that Ycg1 is indeed less stable in the mutant compared to the wild type. The difference between the slopes of the lines depicting the Ycg1 level was also significant, as calculated using GraphPad Prism 8.0.1 (*p* = 0.03). We also tested whether the Ycg1 stability was also impaired in the *ctf19Δ* cells in mitosis. For that, we first arrested both the wild type and *ctf19Δ* cells in metaphase by nocodazole (Fig. S9A) and then added CHX in the media to monitor Ycg1 stability. We found no difference in the Ycg1 stability in the wild type and *ctf19Δ* cells (Fig. S9B, C), indicating that Ctf19 influences the stability specifically in meiosis. From our results, we conclude that higher stability of Cdc5 causes accumulation of the protein in *ctf19*Δ meiotic cells, which results in hyper-phosphorylation of condensin. This hyper-phosphorylated condensin is less stable and possibly why its association with the chromatin decreases (Fig. 5, S10).

### Misregulation of Cdc5 influences chromatin-condensin association

If excess Cdc5 is responsible for reduced chromatin-condensin association in *ctf19*Δ meiotic cells, then overexpression of Cdc5 in the wild type cells should emulate that, whereas its depletion should rescue the defect in the mutant. The overexpression of Cdc5 in the wild type was achieved by shuffling the endogenous promoter of *CDC5* with the *CUP1* promoter. Since overexpression of Cdc5 leads to multiple Spc42 foci (Song et al., 2000), we used this phenotype to optimize the concentration of copper to be used for normal and overexpression of Cdc5. While the cells grown in 20 μM CuSO_4_ in YPD showed very few cells with >2 Spc42-EGFP foci, the same was increased significantly when the cells were grown in 50 μM or higher concentration of CuSO_4_ (Figure S11). Based on this, we assumed that a concentration lesser than 20 μM and higher than 50 μM of CuSO_4_ can be considered as a normal and overexpression of Cdc5, respectively. We further verified the overexpression in meiosis as a phenotype of metaphase I delay, as reported earlier (Shirk et al., 2011, Li et al., 2014). We see a gradual drop in the percentage of cells with metaphase I spindle from 7 hr to 9 hr (59% to 42%) when the cells were treated with 10 μM CuSO_4_ in SPM. However, the cells grown in 50 μm CuSO_4_ did not show the drop (49% to 46%) suggesting sustained metaphase I delay due to over expression of Cdc5 (Fig S12A, B). The cells were released into SPM either in 10 or in 50 μm CuSO_4_ and were harvested at 7 hr (Fig. 9A), since this time point produced a maximum number of metaphase I cells while treated with 50 μm CuSO_4_ (Fig. S12B). In the harvested cells, when we visualized the level of chromatin-condensin association using chromatin spreads, the normal level of Cdc5 (10 μm CuSO_4_) produced 11 to 13 bright Brn1-6HA foci, while with excess Cdc5 (50 μm CuSO_4_), the number was much lower (∼7) (Fig. 9B, C, S12 C) indicating a defect in the association which is consistent to an earlier report (Li et al., 2014).

**Figure 9.**
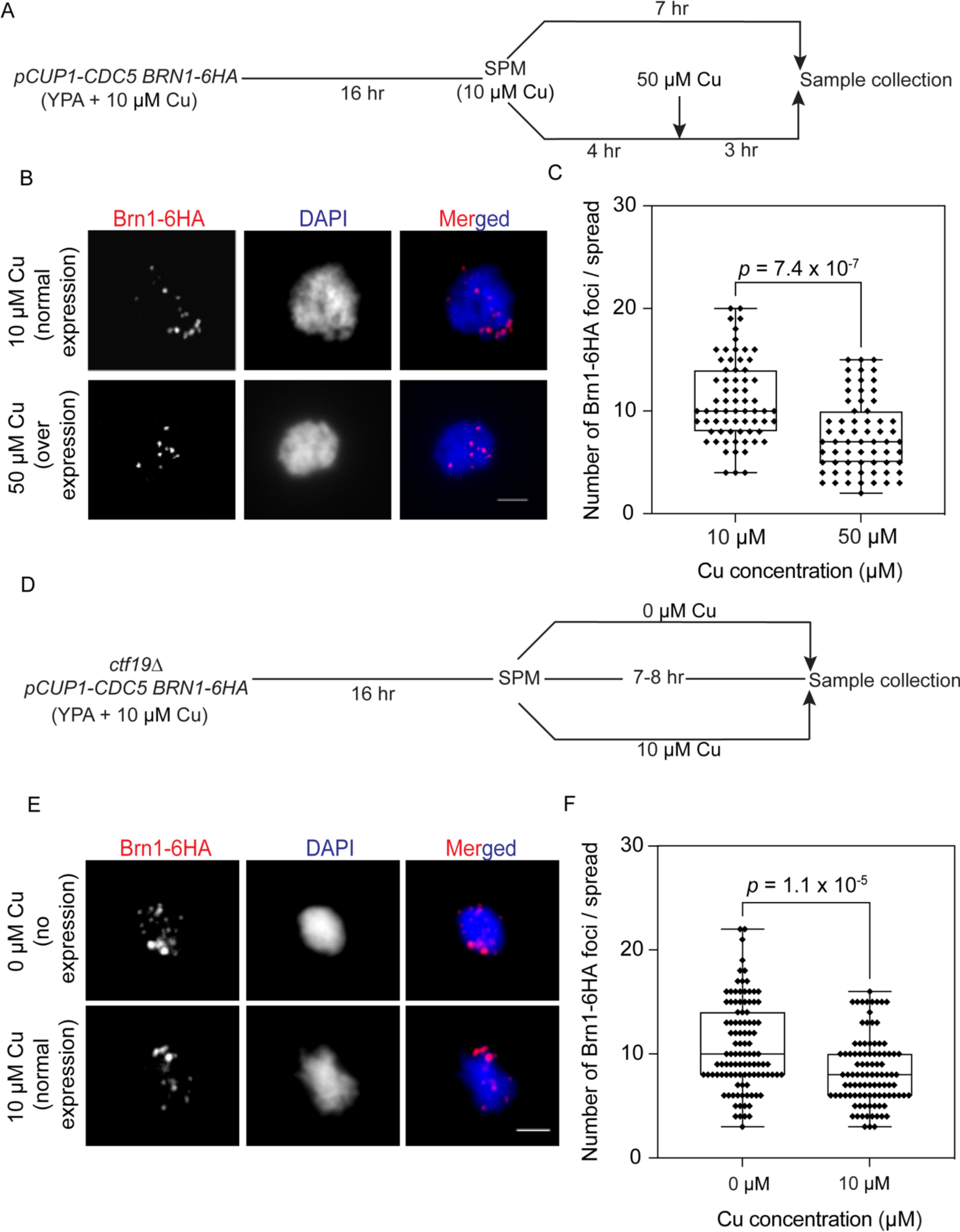
Misregulation of Cdc5 influences chromatin-condensin association. (A) Schematic representation of the experimental strategy. Wild type cells (SGY18007) with homozygous *pCUP1-CDC5* and *BRN1-6HA* alleles were grown in YPA supplemented with 10 μM copper used for normal expression of *CDC5.* The cells were released into SPM supplemented with either 10 μM or 50 μM (used for over expression of *CDC5*) copper, respectively. The samples were harvested at the indicated time point for analysis by chromatin spread assay. (B) Representative images of the chromatin spreads showing chromatin-Brn1-6HA association under normal or overexpression of *CDC5* in the wild type cells. Anti-HA antibody (3F10) was used to detect Brn1-6HA. (C) Box plots showing the number of Brn1-6HA foci per nuclear spread of the wild type cells. N = 60 from two independent experiments. (D, E, F) Same as in A, B and C, respectively except *ctf19Δ* cells (SGY18015) and 0 μM copper was used instead of wild type cells and 50 μM copper, respectively. For F, N = 80 from two independent experiments. Scale bar = 2 μM. The statistical significance *p* value was estimated by the two-tailed student’s t-test for the mean.

Next, to examine if the defect in chromatin-condensin association in the *ctf19Δ* meiotic cells can be rescued by depleting Cdc5, we shuffled the *CDC5* promoter with *CUP1* promoter in those cells and followed the above strategy except used 0 μm CuSO_4_ instead of 50 μm (Fig. 9D). We verified that at 7-8 hrs of SPM release, maximum number of metaphase I cells were present (Fig. S12D, E). With normal level of Cdc5 (10 μm CuSO_4_), we observed around 6 Brn1-6HA foci, while with no Cdc5 expression (0 μm CuSO_4_), we could see a significantly higher number of Brn1-6HA foci (∼10) (Fig. 9E, F, S12 F) indicating a clear amelioration of the defect. Taken together, the above results suggest that the observed meiosis specific condensation defect in the absence of Ctf19 is indeed due to accumulation of Cdc5 in the cells.

## Discussion

For the last few decades, the kinetochore-centromere has emerged as a controlling hub for various events of chromosome segregation viz., kinetochore-microtubule attachment, early replication of centromeres, pericentromeric cohesin enrichment, homolog pairing, and chromosome condensation. In budding yeast, the essential kinetochore proteins have been shown to initiate chromosome condensation in *cis* which is believed to extend distal to the centromeres through Sgo1 and a histone deacetylase, Hst2 (Kruitwagen et al., 2018). On the other hand, through a series of studies, we and others have demonstrated that the organizational assembly of the kinetochore complex differs between mitosis and meiosis and possibly due to this difference it was found that the mitotically non-essential kinetochore proteins play significant roles in meiosis; consequently, in their absence, meiosis is very poor but not mitosis (Mehta et al., 2014, Agarwal et al., 2015, Borek et al., 2021). In this work, we demonstrate that the mitotically dispensable kinetochore proteins contribute to achieve a higher chromatin condensation in meiosis through regulating the activity of Cdc5 polo-like kinase.

### Chromosome condensation: mitosis vs meiosis

Regardless of the mode of the cell cycle, chromosomes need to be condensed as they embark into partitioning. However, there are fundamental differences in how the chromosomes behave as they enter into mitosis or meiosis (meiosis I, in particular). One of the major differences is the pairing of the homolog in meiosis which is believed to start at the centromeres by ‘centromere coupling’ (Tsubouchi et al., 2005) and proceeds towards the telomeres in a zipper-like fashion. It is obvious that finding the right partners and the completion of lengthwise pairing will be more efficient if the units undergoing pairing are shorter and more condensed. Moreover, the post-pairing events, including chiasmata formation and resolution of the chiasmata by ‘terminalization’ of the cross-over point, are also presumably favored by more condensed chromosomes. Notably, in pachytene during meiosis I, the paired homologs are individually visible as thread-like structures by whole chromatin staining using DAPI even in small yeast nuclear spreads. However, although the homologs never pair in mitosis, they never appear as thread-like structures even with half the DAPI intensity of the pachytene spreads. By using two different methods of in vivo condensation assays, we observed that in budding yeast meiotic chromosomes are indeed more condensed than the equivalent mitotic stage chromosomes (Fig. 1, 4C, S3A-B). However, since we observed a greater decrease in intensity of GFP foci and distance between them, as readout for chromatin’s local compaction and global contraction, respectively, in meiosis, both nucleosome-nucleosome interaction and condensin function may culminate into higher meiosis-specific condensation. Evidence of more chromosome condensation in meiosis is available in nature. For instance, in Rye (*Secale cereale*) it is reported that chromosomes are compacted in meiosis by a factor of 1.7 in length and 1.3 in diameter more compared to mitosis (Zoller et al., 2004). Notably, we found ∼1.5 times more condensation in meiosis compared to mitosis by in vivo assay in budding yeast (Fig. 1D) indicating a similar mechanism across eukaryotes might exist to achieve more chromosome condensation in meiosis.

### Kinetochore proteins controlling chromosome condensation

Chromosome condensation can be divided into two stages viz., establishment and maintenance, which are regulated by kinases. As the cells enter into the cell cycle, the condensin becomes active and condensation is established through CDKs (Kimura et al., 1998, Sutani et al., 1999). After the metaphase to anaphase transition, as the CDK level starts declining, aurora B kinase and polo-like kinases phosphorylate condensin subunits and thus maintain the condensation (Lavoie et al., 2004, Takemoto et al., 2007, St. Pierre et al., 2009, Walters et al., 2014, Kruitwagen et al., 2018). In budding yeast mitosis, the essential inner kinetochore proteins promote condensation at the centromeres/pericentromeres by recruiting Ipl1 (aurora B) and Cdc5 (polo-like) kinases at the centromeres (Lavoie et al., 2004, St. Pierre et al., 2009). In contrast, the outer essential and the inner non-essential kinetochore proteins were found to be dispensable for condensation (Nakazawa et al., 2008, Kruitwagen et al., 2018) possibly because they do not regulate chromatin localization of Ipl1 and Cdc5. However, given our earlier study reporting meiosis specific reduction in chromatin-condensin association in absence of the non-essential kinetochore protein Ctf19 (Mehta et al., 2014), in this study we show that the effect is specific to the lack of Ctf19c proteins and it is the establishment of the association which is compromised.

Since our and other studies showed reduced pericentric cohesin enrichment in *ctf19Δ* cells (Mehta et al., 2014, Fernius and Marston 2009), it can be argued that the observed reduction in chromatin-condensin association in *ctf19Δ* meiotic cells is merely due to pericentric cohesin enrichment defect. This can happen provided i) compared to mitosis, in meiosis, the arm cohesin is sufficiently low due to the meiosis-specific prophase-like pathway of cohesin removal (Challa et al., 2019), which becomes aggravated due to simultaneous loss of cohesin from the preicentromeres and ii) the presence of cohesin must promote condensin binding. But we categorically negate the above possibilities because i) half of the arm cohesin still remains after cohesin removal by a prophase-like pathway (Challa et al., 2019) although earlier works failed to detect any significant difference in arm cohesin between mitosis and meiosis (Glynn et al., 2004, Fernius and Marston 2009) and ii) it was shown that cohesin does not influence condensin binding using biochemical and cell biological assays (Lamothe et al., 2020, Lavoie et al., 2002).

Condensation distal from the centromeres can be influenced by non-essential kinetochore proteins through Bub1 and DDK kinases mediated recruitment of Sgo1 (Fernius et al., 2007, Peplowska et al., 2014) and cohesin (Lavoie et al., 2004, Natsume et al., 2013, Hinshaw et al., 2015), respectively, at the pericentromeres which is then believed to extend towards the telomeres through functions of Sgo1 and Hst2 (Kruitwagen et al., 2018). Therefore, we tested whether Sgo1’s centromeric localization and DDK activity are further down in meiosis in *ctf19Δ* cells to explain meiosis-specific condensation defects in these cells. We failed to see any effect on condensation in DDK-compromised cells (Fig. S6A, B and C). However, a higher reduction of Sgo1 level at the pericentromeres in meiosis than in mitosis in *ctf19Δ* cells (Fig. 6) argued this as a possible cause of meiosis-specific condensation defect in these cells. Additionally, we observed that the centromeric association of Ipl1 kinase remains unaffected in *ctf19Δ* cells, indicating that Ctf19 does not play a regulatory role in condensation through the Ipl1-mediated pathway (Fig. S7).

Further studies surprisingly revealed that it is the higher accumulation of Cdc5 in the *ctf19Δ* meiotic cells that perturbs chromatin-condensin association as discussed below.

### Interaction of kinetochore with polo-like kinase

In budding yeast, the polo-like kinase Cdc5 is targeted to the centromeres through interaction with the kinetochore proteins in mitosis (Snead et al., 2007, Mishra et al., 2019). It is also shown that pericentromeric cohesin enriches Cdc5 at the centromeres which in turn promotes cohesin removal (Mishra et al., 2016). Therefore, as *ctf19Δ* cells show reduced pericentric cohesin enrichment (Fernius et al., 2009, Mehta et al., 2014), it is expected that the association of Cdc5 with the chromatin would be less, and that is what we observed (Fig. 7C). But more reduction of Cdc5-chromatin association in meiosis than in mitosis (Fig. 7D) may account for a greater effect of pericentric cohesin loss in meiosis in *ctf19Δ* cells. Earlier, it was reported that the Cdc5 dislodged from the chromatin due to global cohesin inactivation, hyper-phosphorylates condensin which remains bound to the chromatin and is stimulated to misfold the chromatin into a decondensed state (Lamothe et al., 2020). We also noticed hyper-phosphorylation of condensin (Fig. 8A, B), but in contrast observed a huge drop in chromatin-condensin-association in the spreads made from *ctf19Δ* cells (Fig. 5, S4A, S10), possibly because the hyper-phosphorylated condensin generated in our conditions is less stable (Fig. 8C). Notably, while Lamothe et al., 2020, could not find the difference in total Cdc5 level between wild type and cohesin-depleted cells, we observed a substantial increase in Cdc5 level in *ctf19Δ* cells compared to the wild type. That might have caused a different type of hyper-phosphorylation of condensin in our cells compared to their conditions, leading to the dissociation of condensin from the chromatin. Therefore, we conclude that in *ctf19Δ* cells, the meiosis-specific altered behavior of Cdc5 is not only due to pericentric cohesin reduction but also due to its higher stability (Fig. 7B). Currently, it is not known that how the loss of Ctf19 leads to a higher stability of Cdc5 in meiosis. Cdc5 has meiosis-specific roles including degradation of Pds1 in meiosis I (Attner et al., 2013), mono-orientation of sister chromatids by regulating the association of Mam1 protein with kinetochores, step-wise cohesin removal (Lee and Amon, 2003) and dissolution of homolog linkage (Yu and Koshland, 2005). Moreover, unlike mitosis, in meiosis, Cdc5 is degraded by APC associated with meiosis-specific co-factor Ama1 (Okaz et al., 2012). Therefore, a meiosis-specific protein-protein interaction involving Cdc5 is expected. In this context, given a difference in the assembly of kinetochore exists between mitosis and meiosis (Mehta et al., 2014, Borek et al., 2021), the loss of Ctf19 and possibly other non-essential proteins of Ctf19c might create a condition that disfavors normal degradation of Cdc5. One possibility is that in the *ctf19*Δ mutant cells the activity of Mnd2, an antagonist to Ama1 (Oelschlaegel et al., 2005), increases as Mnd2 is believed to physically interact with Ctf19 (Wong et al., 2007). It will be interesting to investigate the status of Mnd2 and Ama1 in the *ctf19*Δ mutant.

Overall, in this work, we reveal that the chromosomes are more condensed during meiosis than in mitosis which has implications for understanding the complex meiotic chromosome behavior in mammals where the gametogenesis is largely asymmetric and mechanistically varies profoundly between the sexes. In animal cells, there is no evidence of kinetochore inactivation affecting chromosome condensation in meiosis. Since we observed that the non-essential proteins affect condensation in meiosis but not in mitosis, it would be intriguing to examine the condensation status in animal meiotic cells harboring mutations in kinetochore. Finally, the interaction between kinetochore integrity and the functionality of polo-like kinase (Plk) can be explored in animal cells as improper polo-like kinase activity has been linked to aneuploidy and different disease states in humans viz., elevated Plk level has been reported in several types of cancers and neurological disorders (Takai et al., 2005, Ackermann et al., 2011, Korns et al., 2022).

### Material and methods Yeast strains

All the strains utilized in this work were derived from the SK1 background and are mentioned in Table S1. Chromosome V was fluorescently marked near *CEN V* and *TEL V* using [TetO]_n_ –[TetR-GFP] system in strains expressing TetR-GFP. An array of 224 copies of the TetO sequence was inserted at locus 1.4 kb away from *CEN V* through homologous recombination (Michaelis et al., 1997, Tanaka et al., 2000). Similarly, [TetO]_448_ array was inserted near *TEL V* between *BMH1* and *PDA1* genes (Kumar et al., 2021). To perform gene deletion, promoter shuffling, and C-terminal tagging of proteins, a PCR-based method was used (Wach, 1996, Longtine et al.,1998, Janke et al., 2004). Relevant plasmid-borne cassettes procured from Euroscarf were employed for these techniques. Yeast transformation was performed as described previously (Guldener et al., 1996).

The cells were synchronized in the G1 phase in the YPA medium and then released in YPD and SPM for mitotic and meiotic sample collection, respectively. For the mitotic pre-S phase, the cells were harvested at 45 mins post-release in YPD which corresponds with the start of Cln2 expression. The cells were identified microscopically by unbudded cells with one SPB focus. For metaphase, the cells were harvested after 1.5 hrs post-release in YPD and distinguished microscopically by large-budded cells with two SPB foci in the mother bud. For meiosis, the cells were harvested at 1 hr and 6 hrs post-release in SPM for pre-S phase and metaphase I cell, respectively.

### Meiotic synchronization

Meiotic induction was carried out as described elsewhere (Cha et al., 2000). Typically, the diploid strain of interest was streaked onto a YPD plate (1% yeast extract, 2% peptone, 2% dextrose, 2% agar) for a single colony followed by 48 hrs of incubation. A single colony was then inoculated into 5 ml of YPD broth and left to incubate overnight at 30°C for ∼15 hrs. Subsequently, cells from the overnight grown culture were transferred to a 50 ml YPA medium (1% yeast extract, 2% peptone, and 1% potassium acetate) at 0.2 OD_600_ and grown till ∼1.4-1.6 OD_600_ which took approximately 13-15 hrs. 1 ml of YPA culture was taken, the cells were diluted appropriately and subjected to sonication at 50% amplitude for 3 secs and observed under microscope for G1 arrest. Once over 85% of G1 arrest was achieved, the cells were washed with sterile distilled water and resuspended in 50 ml of sporulation media (SPM; 0.3% potassium acetate and 0.02% raffinose). Meiotic synchronization was performed at 30°C with continuous shaking at 200 RPM. Samples were harvested from the SPM at specified time intervals for further experimentation. For metaphase I arrest, the cells harboring *CDC20* promoter shuffled with mitotic *CLB2* promoter (Lee and Amon, 2003), were harvested at 8 hrs and 10 hrs for wild type and *ctf19*Δ mutant, respectively.

### The arrest of yeast strains at mitotic metaphase

#### Using the auxin-induced degron system

Mitotic metaphase arrest was achieved by degrading Cdc20 using the AID degron (Auxin-Inducible Degron) system (Nishimura et al., 2009). In cells expressing *OsTIR1* from the *ADH1* promoter, the AID coding sequence was C-terminally fused to *CDC20*. Cells were grown in YPD from 0.2 to 0.5 OD_600_ before auxin (3-Indoleacetic acid, Sigma) was added at 1.5 mM concentration and the culture was incubated for a further 2 hrs at 30⁰C. Metaphase arrest was confirmed by DAPI staining (more than 85% large budded cells with undivided DAPI).

#### Using nocodazole, a microtubule depolymerizing drug

The cells were inoculated in YPD at 0.2 OD_600_ and grown till 0.5-0.6 OD_600_ before nocodazole (20 µg/ml, Sigma) was added to the culture which was incubated for a further 2 hrs under shaking conditions at 30⁰C. Metaphase arrest was confirmed by DAPI staining.

### Fluorescence microscopy

The cells were grown in YPD or SPM medium for mitotic or meiotic culture, respectively. The cells were harvested and fixed with 4% formaldehyde for 5 mins and then washed twice with 0.1 M phosphate buffer before imaging. All fluorescence images were acquired with a Zeiss Axio Observer.Z1 (63X objective, NA = 1.40; Carl Zeiss Micro-Imaging Inc.) equipped with Axiocam camera. The images were processed and analyzed using Zeiss Zen 3.1 software. Image J and Imaris (8.0.2) software were used for intensity and distance measurements, respectively. For all the box plots for the intensity and distance measurements, mean values were compared across the samples.

### Intensity measurement

For intensity calculation of a fluorescence signal, Z-stacks with maximum intensity were merged. A Region Of Interest (ROI) was drawn around the signal of interest, and the integrated density was determined to get the ‘signal intensity’. ROI of the same size was put elsewhere in the nucleus for background signal and integrated density was determined to get the ‘background intensity’. Each signal intensity value was normalized with the background intensity values to get the ‘normalized signal intensity’ which was used to plot the graphs. For *CEN V-*GFP intensity comparison between pre-S phase cells and metaphase I cells in meiosis, the normalized signal intensity obtained from metaphase I cells was divided by 2 to obtain intensity per *CEN V*-GFP as at this stage two sister centromeres are glued together by cohesin and two sister *CEN V*-GFP dots appeared as a single dot (Fig S13).

### Second Harmonic Generation (SHG) microscopy and analysis

The sample preparation for SHG microscopy was followed as mentioned earlier (Yamin et al., 2020; 2022). Briefly, the cells were harvested and fixed in 4% paraformaldehyde for 20 mins at RT; they were washed with 0.1 M phosphate buffer and stained with DAPI (Invitrogen, #D1306) for 20 mins in dark. The cells were subsequently placed on a polylysine coated slide and allowed to adhere to the slide. After washing the immobilized cells with PBS, mounting solution was added and the slide was covered with a coverslip. Cells were then analyzed by a Zeiss LSM780 inverted Confocal Microscope, with multi-photon Chameleon vision II Laser (laser power > 3 W, pulse width and repetition rate at default settings, tuning range 690–1040 nm); Lambda—PMT—identifying all the wavelength spectrums. The slides were excited by 768 nm and emission was collected at 414-628 nm. The quantification was carried out using ImageJ software as mentioned earlier (Yamin et al., 2020; 2022).

### Chromatin spread and the analysis

Chromatin spread was performed by following the protocol mentioned elsewhere (Mehta et al., 2014) with certain modifications. Cells were harvested from the mitotic culture of 1.0 OD_600_ and at the required time point for meiotic culture. The cells were washed once with spheroplasting solution (1.2 M sorbitol in 0.1 M phosphate buffer) and harvested by centrifugation at 3000 RPM for 3 mins. The cells were resuspended in 0.5 ml of spheroplasting solution supplemented with 5 µl of β-mercaptoethanol and 12.5 µl of zymolyase 20T (10 mg/ml, MP Biomedicals) and incubated at 30⁰C for 1.5 hrs. Once 80-90% of the cells were spheroplasted (by observing under the microscope), the spheroplasting was stopped by adding 1 ml of ice-cold stop solution (0.1 M MES - pH 6.4, 1 mM EDTA, 0.5 mM MgCl_2_, 1 M Sorbitol). The spheroplasts were centrifuged at 2000 RPM for 3 mins at 4 ⁰C and resuspended gently in 125 µl of ice-cold stop solution. 60 µl of spheroplasted cells were put on acid-washed glass slides followed by the addition of 40 µl of fixative solution (4% paraformaldehyde, 3.4% sucrose, 10 N NaOH) and 80 µl of 1% lipsol (LIP Equipment and services) over it. After gentle mixing 80 µl of fixative solution was again added to the slides. The content on the slides was mixed and spread gently using a pipette tip and kept at room temperature overnight for drying. The slides were then washed with 2 ml of 0.4% photo flow-200 (Kodak) and then submerged in 1X PBS for 15 mins. The excess liquid was removed and 100 µl of blocking solution (10 mg/ml BSA in 1X PBS + 5% Skim milk) was immediately added over the fixed nuclear spreads which were then covered with coverslips and incubated at room temperature in a humid chamber for 20 mins. The coverslips were removed and 100 µl of primary antibody (1:200) was added to each slide and incubated at room temperature for 60 mins. The primary antibody was rinsed off by submerging the slides 3 times in 1X PBS for 5 mins each and 100 µl of secondary antibody (1:200) was added followed by incubating at room temperature for 60 mins. The slides were again washed in the same way followed by incubation with 100 µl of DAPI (1 µg/ml, Invitrogen) for 30 mins. After PBS wash for 5 mins, 100 µl of mounting solution (90% glycerol supplemented with 1 mg/ml p-phenylenediamine) was added over the slides followed by a clean coverslip which was sealed with transparent nail paint. The primary antibodies used: rat anti-HA (3F10, Roche), mouse anti-HA (12CA5, Roche) and rat anti-Tubulin (MCA78G, Serotec). The secondary antibodies used: Rhodamine (TRITC)-conjugated goat anti-Rat (Jackson) and Alexa Fluor 488 conjugated goat anti-Mouse IgG (Jackson).

For calculation of the number of fluorescence foci associated with each chromatin spread, all the Z-stacks with the signals were first merged and then the file was exported to Imaris (8.0.2) software. The images were visualized using ‘3-D view’ tool of the software. Subsequently, the ‘Spot’ tool in the algorithm setting of the software was used to mark the different foci present in the chromatin spreads. The background fluorescence was subtracted using an automated ‘subtraction’ tool, and the number of foci visible for each spread was scored.

### Indirect immunofluorescence

Immunofluorescence was performed, as mentioned previously (Mittal et al., 2020; Shah et al., 2023). Briefly, cells from meiotic culture were harvested at indicated time points and fixed with 4% formaldehyde at room temperature (RT) for 15 mins. The fixed cells were washed once with PBS and spheroplasted in spheroplasting solution (1.2 M sorbitol, 0.1 M phosphate buffer pH 7.5), in the presence of Zymolyase 20T (MP Biomedicals, #32092) (10 mg/ml) and 25 mM β-mercaptoethanol or 1 hr at 30°C. The spheroplasts were mounted on a polylysine coated slide. The cells were then flattened by immersing the slide in -20°C methanol for 5 mins and in acetone for 30 s. The slides were dried for 1-2 mins before blocking using 5% skim milk solution prepared in dilution buffer (10 mg/ml BSA in PBS). The cells were incubated with appropriate primary and secondary antibodies for 1 hr at RT. Following several washes with PBS, samples were incubated with DAPI (Invitrogen, #D1306) (1 μg/ml) solution for 20 mins in dark. Finally, the glass slides were sealed with coverslips for imaging after spreading the mounting solution (1 mg/ml phenylenediamine in 90% glycerol). The antibodies and their dilutions (in dilution buffer) were mentioned as follows. Primary antibodies rat anti-tubulin (Serotec, MCA78G) at 1:5000. Secondary antibodies: (TRITC)-labeled goat anti-rat (Jackson, #115485166).

### ChIP (Chromatin immunoprecipitation)

Chromatin immunoprecipitation was performed as described previously (Makrantoni et al., 2019) with certain modifications. Chromatin was cross-linked with 1% formaldehyde (30 mins for Brn1- 6HA, 1 hr for Sgo1-9Myc, 2 hrs for Ipl1-6HA) at 25℃ with shaking of the culture at 100 RPM, followed by the addition of glycine at a final concentration of 125 mM to quench the crosslinking. Cells were harvested and washed twice with 10 ml ice-cold TBS buffer and once with 10 ml 1X FA lysis buffer (50 mM Hepes–KOH pH 7.5, 150 mM NaCl, 1 mM EDTA, 1% Triton X-100, 0.1% Na-deoxycholate) supplemented with 0.1 % SDS. The cell pellet was resuspended in 500 μl of ice-cold 1X FA lysis buffer supplemented with 0.5% SDS, 1 mM PMSF and 1X PIC (Protease Inhibitor Cocktail, Roche). An equal volume of glass beads was added and cells were lysed in a mini bead beater (BIOSPEC, India) for 9 cycles (1 min ON, 1 min OFF on ice). The lysate was collected in a pre-chilled 1.5 ml eppendorf tube and centrifuged at 16,000 x g for 15 min at 4°C. The supernatant was removed and the pellet was resuspended in 1 ml ice-cold 1X FA lysis buffer supplemented with 0.1% SDS, 1 mM PMSF and PIC. The tubes were again centrifuged at 16,000 x g for 15 mins. The supernatant was discarded and the chromatin pellet was resuspended in 300 μl ice-cold 1X FA lysis buffer supplemented with 0.1% SDS, 1 mM PMSF and PIC. The chromatin was then sheared to 300-500 bp using a water bath sonicator (Diagenode SA, Picoruptor, BC 100, LAUDA Germany) for 30 cycles of 30 s ON / 30 s OFF at 4°C. The sonicated sample was then centrifuged at 16,000 x g for 15 mins at 4°C. The supernatant was collected to which 1X FA lysis buffer was again added and re-centrifuged at the same settings. 10 µl of the supernatant was stored as input at -20℃ (Whole cell extract, WCE) and the rest was divided into ‘plus antibody’ (+Ab) and ‘minus antibody’ (-Ab), 600 µl each. Aliquots (+Ab) were incubated with 5 µg antibody [anti- HA (3F10, Roche) or anti-MYC (AB9106, Abcam)] overnight in rotating conditions followed by incubation with 50 µl of 50% Protein A Sepharose-beads (GE Healthcare,17-0780-01) for 2 hrs at 4°C in rotating condition. The beads were washed at room temperature twice with 1 ml of IP wash buffer I (1X FA lysis buffer, 0.1%SDS, 275 mM NaCl), twice with 1 ml of IP wash buffer II (1X FA lysis buffer, 0.1%SDS, 500 mM NaCl), once with 1 ml of IP wash buffer III (10 mM Tris pH 8.0, 250 mM LiCl, 0.5% NP-40, 0.5% sodium deoxycholate, 1 mM EDTA) and once with 1 ml of 1X TE (Tris EDTA). The beads were resuspended in elution buffer (50 mM Tris, pH 8.0, 10 mM EDTA, 1% SDS) and incubated overnight at 65°C for de-crosslinking. De-crosslinked samples were treated with Proteinase K (SRL, India) at 45 °C for 2 hrs followed by P:C:I (Phenol:Chloroform:Isoamyl, Himedia) purification. Purified samples were further used for qPCR analysis using CFX96 Real-Time system (BioRad) and SYBR green dye (BioRad) for detection. Enrichment/Input was calculated by the following equation: % 𝐸𝑛𝑐𝑟𝑖𝑐ℎ𝑚𝑒𝑛𝑡/𝑖𝑛𝑝𝑢𝑡 = 𝐸^(−ΔCt); where, E is the primer efficiency and was calculated by plotting Ct values of serially diluted input with respect to Log dilution. E = (−1)/slope and *ΔCt = Ct _(Test sample)_- (Ct _(Input)_-Log E (Input dilution factor).* Primers used for PCR amplification are given in Table S2. For calculation of percentage decrease of Sgo1-9Myc enrichment at the centromeres in the ChIP assays (Fig 6C and D) following formulae was used: 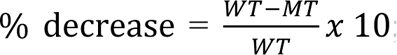; (WT and MT are average enrichment/input values from the wild type and the mutant, respectively).

### Protein isolation and western blotting

Samples for western blotting were prepared by resuspending the cells in 200 µl of 20% TCA. Equal volume of glass beads was added, and bead beating was done using a mini bead beater (BIOSPEC, India) for 5 cycles (1 min ON, 1 min OFF on ice). Supernatant was collected by centrifugating at 2800 RPM for 2 mins. 400 µl of 5% TCA was again added to the beads and supernatant was again collected after centrifugation. The total supernatant obtained (600 µl) was centrifuged at maximum speed (13,500 RPM) for 5 mins to precipitate the protein. The supernatant was removed, and the protein pellet was resuspended in Laemmli buffer and loaded on SDS-PAGE after boiling at 95℃ for 5 mins for further analysis. For phosphatase assay, protocol mentioned elsewhere (Ontoso et al., 2013) was followed with certain modifications. TCA extract in Laemmli buffer was diluted 10-fold in 1X PMP buffer (NEB, Protein Metallo Phosphatases) supplemented with 1 mM Mncl_2_. The diluted sample was treated with 2 µl Lambda protein phosphatase (NEB) and incubated in water bath at 30 ℃ for 4 hrs. As control, a similar diluted extract but without adding phosphatase was incubated at similar conditions. The protein sample was TCA precipitated and resuspended in Laemmli buffer followed by heating at 95℃ for 5 mins. Western blotting was performed using standard procedure. The blots were developed using following antibodies: rat anti-tubulin antibody (MCA78G, Serotec,), mouse anti-HA (12CA5 and 3F10, Roche), HRP-conjugated goat anti-mouse IgG (Jackson), HRP-conjugated goat anti-rat (Jackson). For CHX-chase experiments, the percentage of protein remaining at each time point was calculated by taking the protein level at 0 hr as 100%.

### RNA isolation

Total RNA was extracted from meiotic cell culture of 1.5-2 OD_600_ (∼2.5 x10^7^ cells) using TRIzol® reagent and PureLink® RNA mini kit (Ambion Life Technologies™). The cell pellet was re- suspended in TRIzol® and cell lysis was done using DEPC treated sterile 0.5 mm glass beads. The rest of the protocol was followed as instructed by the supplier (PureLink® RNA mini kit). RNA was eluted in DEPC (Sigma Aldrich Chemicals Pvt. Ltd.) treated sterile water. DNase treatment was done at 37°C for 10 mins using RNAse-free DNase I (Thermo Fisher Scientific) in the buffer supplied with the enzyme. The DNase and salts in the buffer were removed by purification using trizol-based layer separation and pure RNA was recovered from the aqueous layer. To ensure the absence of DNA impurity, end-point PCR was performed.

### Biological replicates and statistical methods

The quantitative data shown in all the figures were obtained from two or three biological replicates. The N values in the figure legends are the total number of cells analyzed from combined replicates of individual assay. Box plots were generated using OriginPro software (OriginLab) and GraphPad Prism 9.0 (Version 9.4.1) software. Statistical significance was estimated using two-tailed student’s t-test for mean with p< .05 considered to be significant.

## Acknowledgement

We acknowledge the central instrumentation facility of IIT Bombay. The SKG laboratory is supported by the Science and Engineering Research Board (SERB), Govt. of India (Grant No. CRG/2020/000444). DT and PA are supported by CSIR fellowships (09/087(0886)/2017-EMR-I), and (09/087(0972)/2019-EMR-I), respectively. AA is supported by JRF fellowship from Department of Biotechnology (DBT), Govt of India (Grant No. BT/PR43050/BRB/10/1992/2021).

## Author contributions

DT, PA and AA performed experiments. DT and SKG designed experiments, analyzed and interpreted the data. DT, PA and SKG wrote the manuscript.

