## Supplementary file 1 for "Non-essential kinetochore proteins contribute to meiotic chromosome condensation through polo-like kinase"

### SUPPLEMENTAL MATERIAL

### SUPPLEMENTAL FIGURES

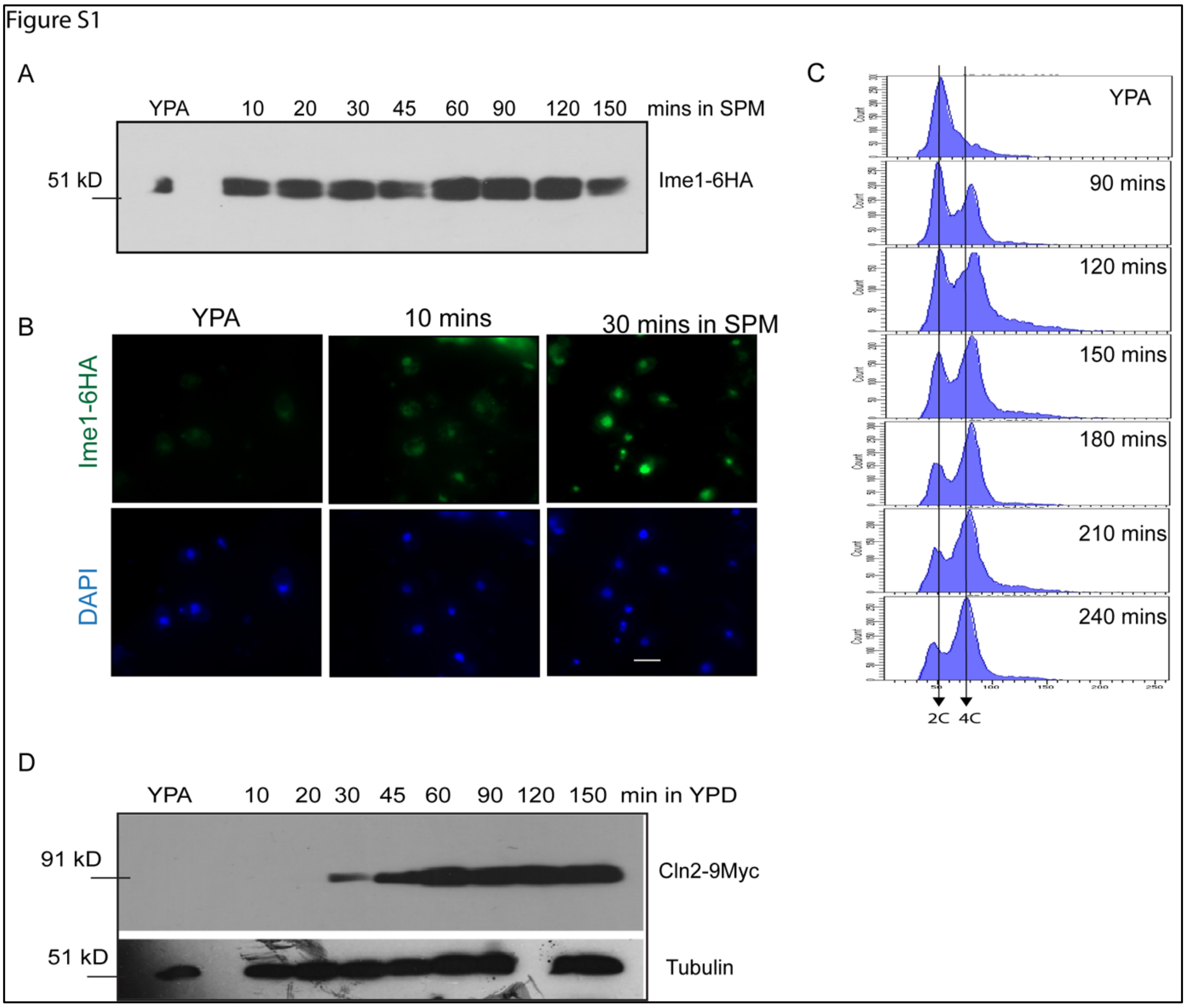

**Figure S1. Determination of the time point to harvest cells residing at the pre-S phase in meiosis.** (A) Ime1-6HA expression in wild type (SGY7098) at different time points after release of G1 cells into SPM (meiosis). Ime1-6HA was detected by anti-HA antibodies (12CA5, 1:2500). (B) Sub-cellular localization of Ime1-6HA was determined in YPA and at indicated time points in SPM by indirect immunofluorescence using anti-HA antibodies (12CA5, 1:200). (C) Premeiotic DNA replication in wild type cells (SGY7098) starts before 90 min in SPM, as evidenced by FACS analysis. The cells were grown in the YPA medium for G1 arrest, washed in water, and resuspended in SPM (time 0). Samples were harvested at the indicated time points for DNA content analysis by FACS. (D) Cln2-9Myc expression in the same strain as above at different time points after the release of G1 cells into YPD (mitosis). Tubulin expression was monitored as a control. Cln2-9Myc and tubulin were detected by anti-Myc (9E10) and anti-tubulin (YOL1/34) antibodies. \

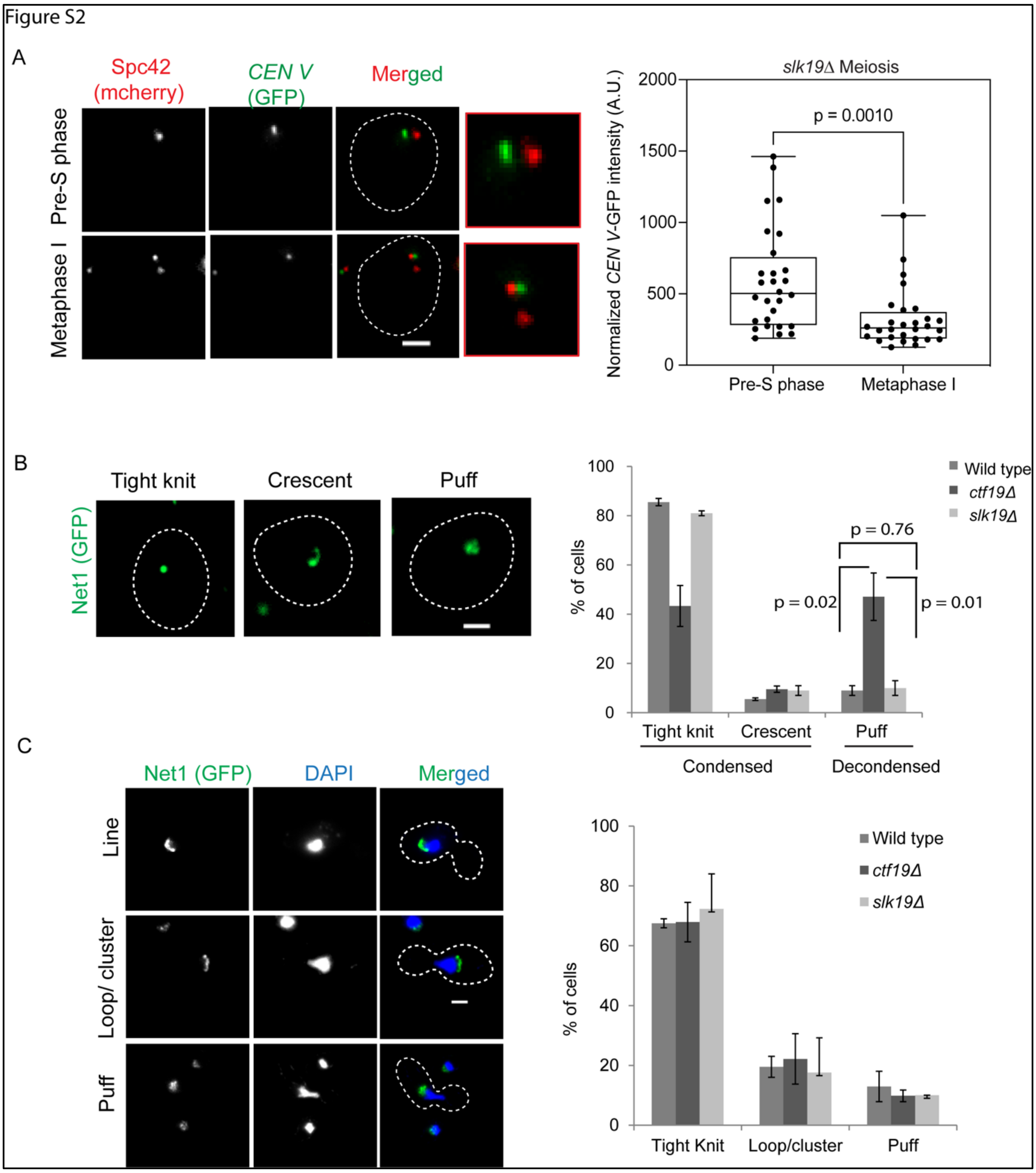

**Figure S2**: **Ctf19 has no role in rDNA condensation in mitosis**. (A) In vivo, chromosome condensation in *slk19*Δ cells (SGY9168) was measured as in Fig. 1 for the indicated stages. The intensity values are graphically represented on the right; the corresponding representative images are shown on the left. N = 25-30 from two independent experiments, scale bar = 2 µm. rDNA condensation was analyzed by visualizing the rDNA-binding protein Net1 tagged with GFP in the wild type (SGY329), *ctf19*Δ (SGY327), and *slk19*Δ cells (SGY 9020) arrested in the meiotic metaphase I (B) stage or mitosis (C) identified by the presence of undivided DAPI staining within the mother compartment of the large budded cells N > 200, scale bar = 2 µm. Error bars represent the standard deviation from the mean values obtained from three independent experiments. For statistical significance, *p* values were estimated by the two-tailed student's t-test for the mean.

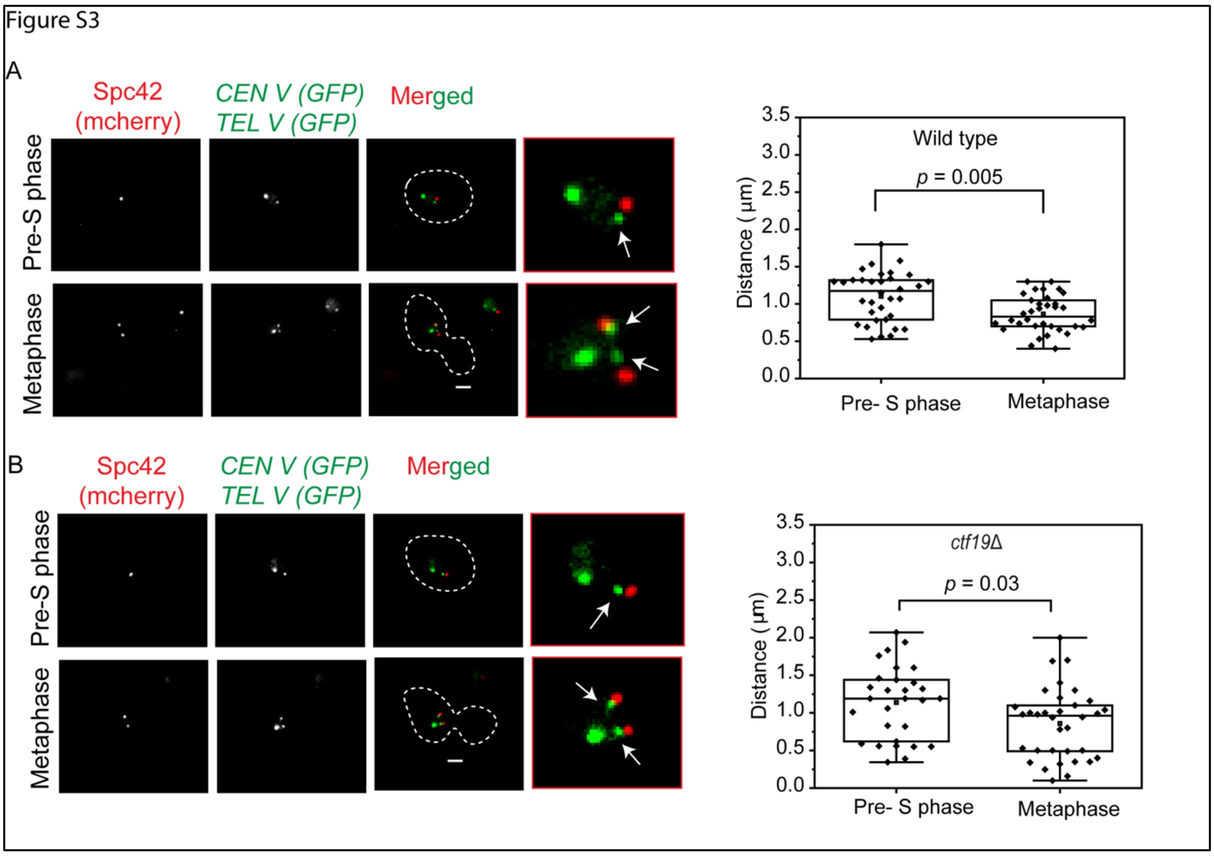

**Figure S3. The condensation along the chromosomal arm is normal in mitosis in the absence of Ctf19.** The 3D distances between the *CEN V*-GFP and *TEL V*-GFP dots measured as described in the materials and methods are shown for the (A) wild type (SGY9092) and (B) *ctf19*Δ (SGY9124) cells harvested from the indicated cell cycle stages in mitosis. The representative images are shown on the left whereas the box plots are shown on the right. N = 27-34 from two independent experiments, scale bar = 2 µm. The statistical significance *p* value was estimated by the two-tailed student's t-test for the mean. Arrows indicate *CEN V* dots.

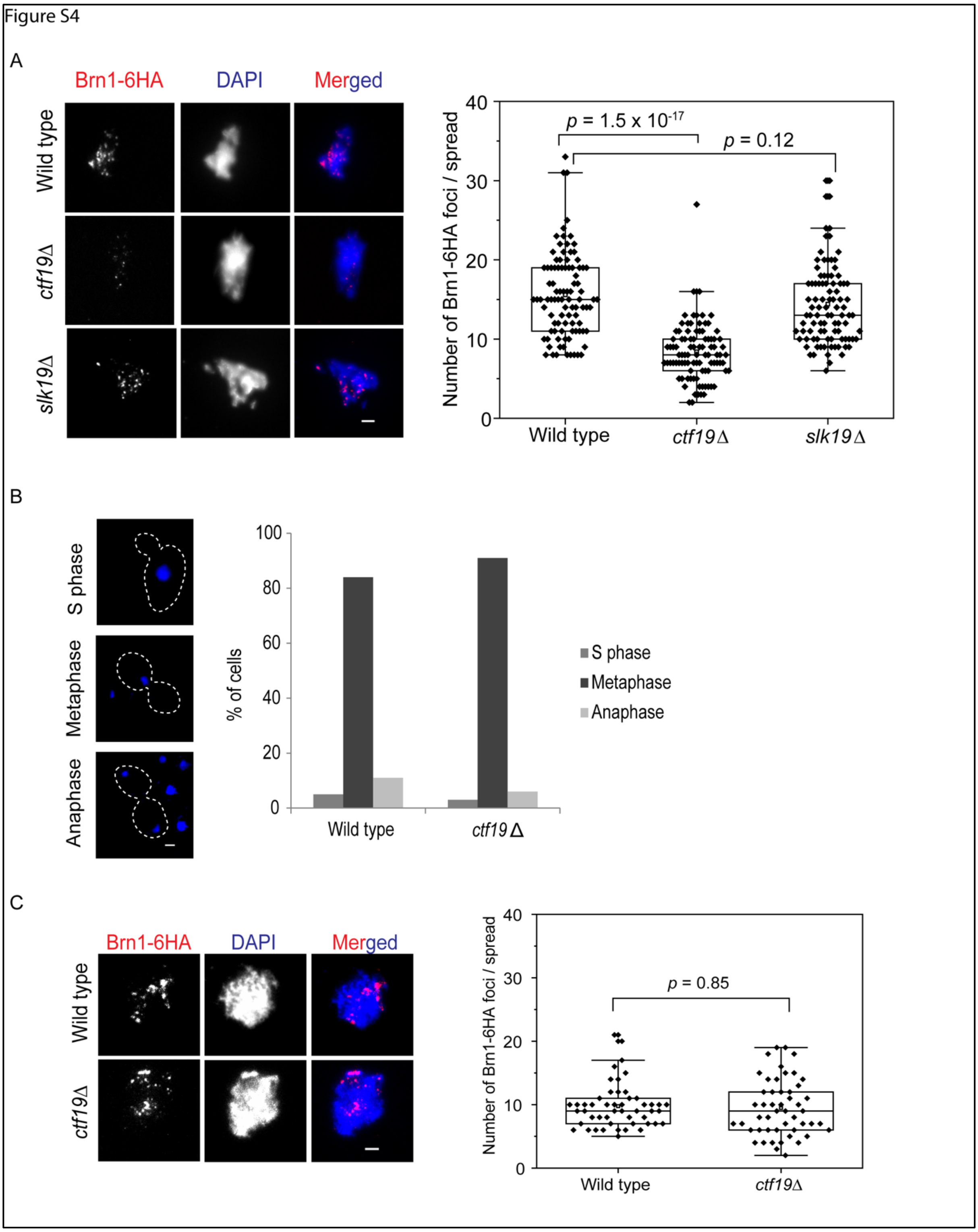

**Figure S4. The loss of Ctf19 affects the association of condensin with chromatin in meiosis but not in mitosis**. (A) Chromatin spreads showing chromatin-condensin association in the metaphase I arrested cells of the wild type (SGY283), *ctf19*Δ (SGY284), and *slk19*Δ (SGY9025) strains harboring homozygous *BRN1-6HA* allele. Anti-HA antibodies (3F10) were used to detect Brn1-6HA. The left panel shows the representative images of the spreads; the right panel represents the box plots showing the number of Brn1-6HA foci per nuclear spread of the indicated strains. N = 80 from three independent experiments, scale bar = 2 µm. (B) The left panel shows the representative images of the wild type (SGY9163) and *ctf19Δ* (SGY9165) strains after treatment with auxin for Cdc20 depletion to achieve metaphase arrest in mitosis (materials and methods); the right panel shows the percentage of the cells in different cell cycle stages. N = 100, scale bar = 5 µm. (C) Chromatin spreads showing chromatin-condensin association in the metaphase arrested strains as in B harboring homozygous *BRN1-6HA* allele. Anti-HA antibodies (3F10) were used to detect Brn1-6HA. The left panel shows the representative images of the spreads; the right panel represents the box plots showing the number of Brn1-6HA foci per nuclear spread of the indicated strains. N = 50 from two independent experiments, scale bar = 2 µm. The statistical significance *p* value was estimated by the two-tailed student's t-test for the mean.

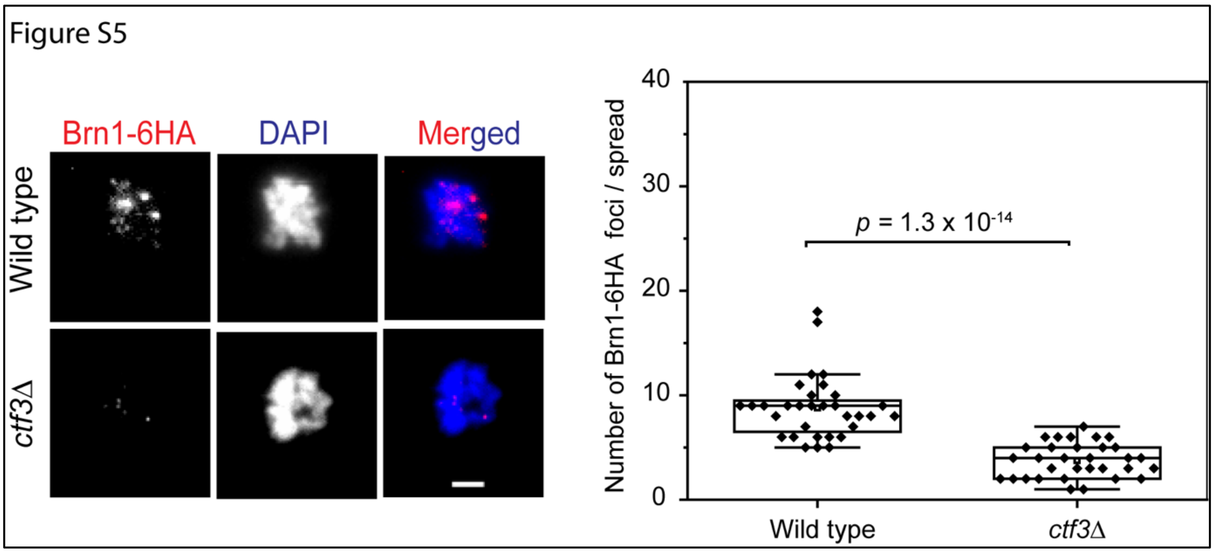

**Figure S5. The loss of Ctf3 causes reduced association of condensin with chromatin in meiosis**. Chromatin spreads showing chromatin-condensin association in the metaphase I arrested cells of the wild type (SGY283) and *ctf3Δ* (SGY9277) strains harboring homozygous *BRN1-6HA* allele. Anti-HA antibodies (3F10) were used to detect Brn1-6HA. The left panel shows the representative images of the spreads; the right panel represents the box plots showing the number of Brn1-6HA foci per nuclear spread of the indicated strains. N=32 from two independent experiments, scale bar = 2 µm. The statistical significance *p* value was estimated by the two-tailed student's t-test for the mean.

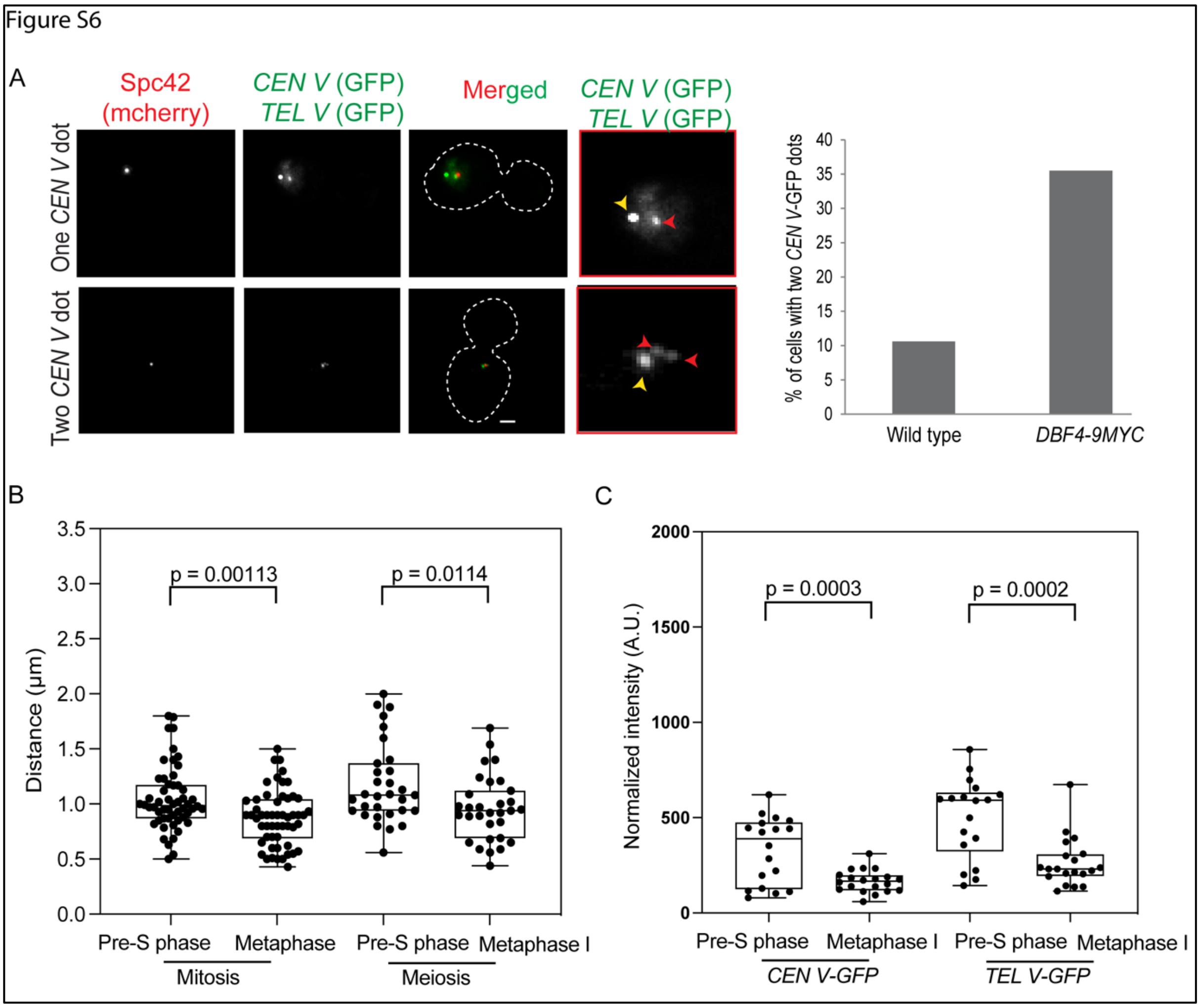

**Figure S6. Chromosome condensation is not affected due to inactivation of DDK kinase.** (A) Sister chromatid cohesion assay in wild type (SGY9092) and *DBF4-9MYC* (SGY9169) cells arrested at metaphase after removing the spindle force using nocodazole. The left panel depicts the representative images of one (cohesed) or two (non-cohesed) *CEN V*-GFP dots whereas the right panel shows the frequency of the latter type. The red and yellow arrowheads indicate *CEN V*-GFP and *TEL V*-GFP dots, respectively. N = 100 from two independent experiments, scale bar = 2 µm. (B) The 3D distances between the *CEN V*-GFP and *TEL V*-GFP dots measured as described in the materials and methods are shown for the *DBF4-9MYC* cells (SGY9169) harvested from the indicated cell cycle stages in mitosis and meiosis. N = 54 (mitosis) and 30 (meiosis) from two independent experiments. (C) The *CEN V*-GFP and *TEL V*-GFP intensities were measured in the *DBF4-9MYC* cells (SGY9169) from the indicated cell cycle stages in meiosis. N = 18-20 from two independent experiments. The GFP intensities were normalized with background intensity. The statistical significance *p* value was estimated by a two-tailed student's t-test for the mean.

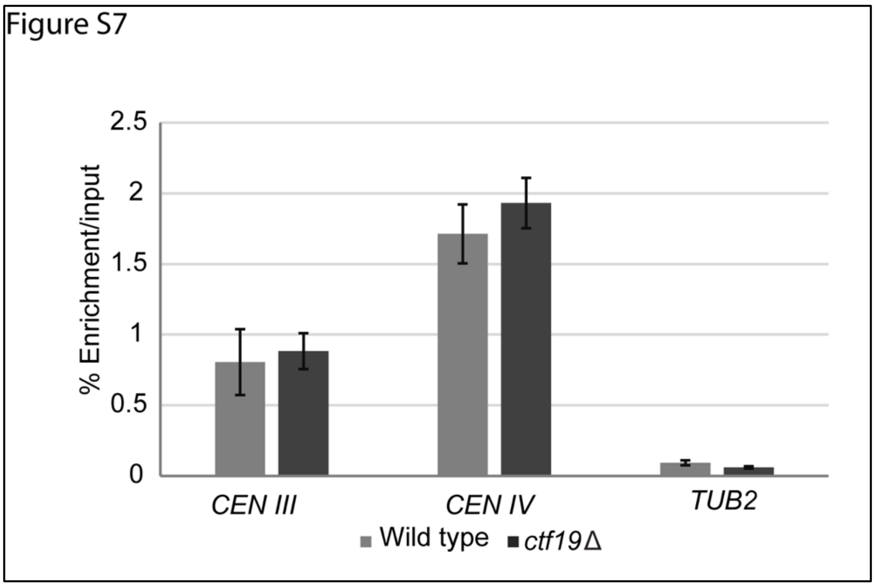

**Figure S7. Loss of Ctf19 does not affect centromeric localization of Ipl1**. ChIP assay using anti-HA (3F10) antibodies for quantifying the association of Ipl-6HA with the indicated chromosomal loci in the wild type (SGY9225) and *ctf19Δ* (SGY9228) cells arrested at metaphase I. The graph represents % enrichment per input obtained by qPCR analysis. Error bars represent the standard deviation from the mean values obtained from two independent experiments.

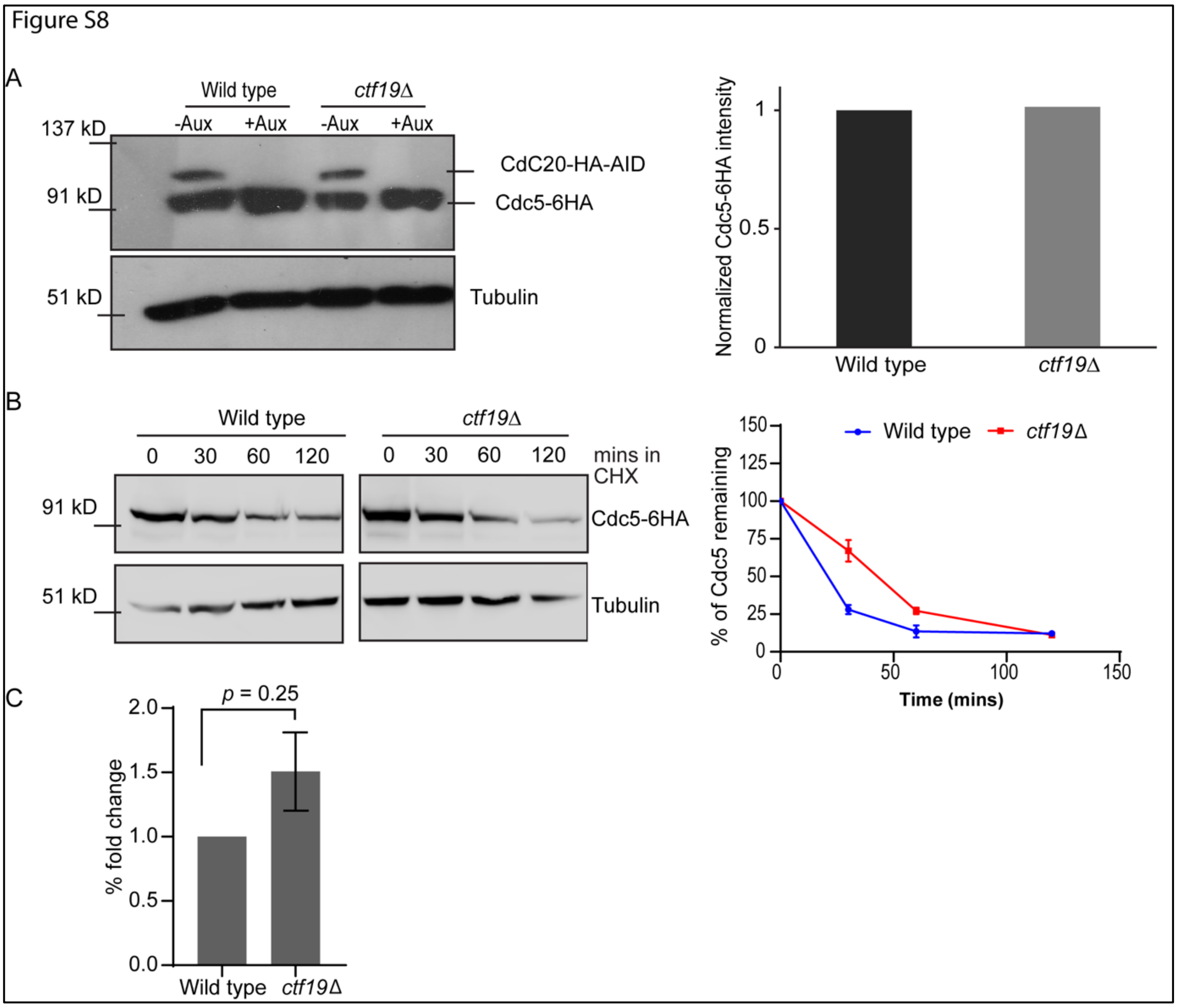

**Figure S8. The level of Cdc5 is similar in the presence or absence of Ctf19 in mitosis.** Proteins were extracted from the wild type (SGY9210) and *ctf19*Δ (SGY9208) cells harboring *CDC5-6HA CDC20-6HA-AID* arrested at metaphase by depletion of Cdc20 using the AID degron system. (A) Left, the western blots show the indicated bands developed using anti-HA and anti-tubulin antibodies. The cells were harvested 2 hrs before (-Aux) and 2 hrs after (+Aux) auxin addition in the media. Right, the intensity of each Cdc5-6HA band was calculated using ImageJ software and normalized to the tubulin band intensity. (B) Cdc5-6HA stability in wild type (SGY9139) and *ctf19*Δ (SGY9140) cells. The cells at the mid-log stage were treated with CHX (t = 0). Left, the protein level was analyzed by western blotting using anti-HA and anti-tubulin antibodies in the cells harvested at indicated time points. Right, quantification of Cdc5-6HA bands normalized with the tubulin bands using ImageJ software at indicated time points after CHX treatment. Error bars represent the standard deviation from the mean values obtained from two independent experiments. (C) Quantification of *CDC5* transcript using qPCR shows no significant change in the Cdc5 expression between wild type (SGY9152) and *ctf19*Δ (SGY9148) cells arrested at metaphase I. Error bars represent the standard deviation from the mean values obtained from two independent experiments.

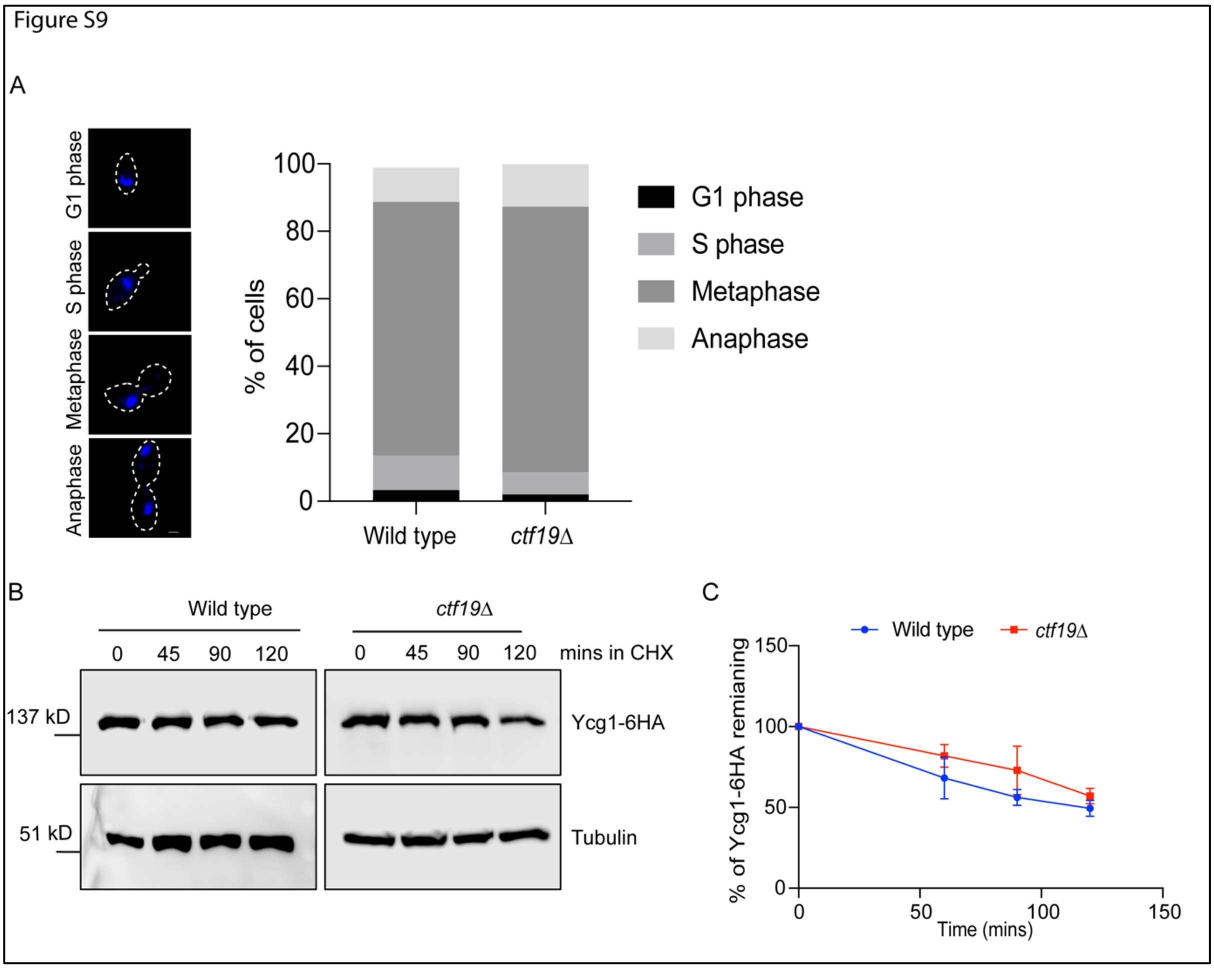

**Figure S9. Loss of Ctf19 does not affect Ycg1 stability in mitosis.** (A) The left panel shows the representative images of the cells at different stages of the cell cycle after treating them with nocodazole for 120 mins. The right panel shows the percentage of the indicated cells for both wild type (SGY9232) and *ctf19*Δ (SGY9233). N = 100, scale bar = 2 μm. (B) Ycg1-6HA stability in wild type and *ctf19*Δ cells. The cells were treated with CHX (t = 0) following mitotic metaphase arrest by nocodazole for the wild type and *ctf19Δ*, respectively. The protein extracts were resolved on 10% SDS-PAGE gel and the protein level was analyzed by western blotting using anti-HA (12CA5) and anti-tubulin (YOL1/34) antibodies in the cells harvested at indicated time points. (C) Quantification of Ycg1-6HA bands normalized with the tubulin bands using ImageJ software at indicated time points after CHX treatment. Error bars represent the standard deviation from the mean values obtained from two independent experiments.

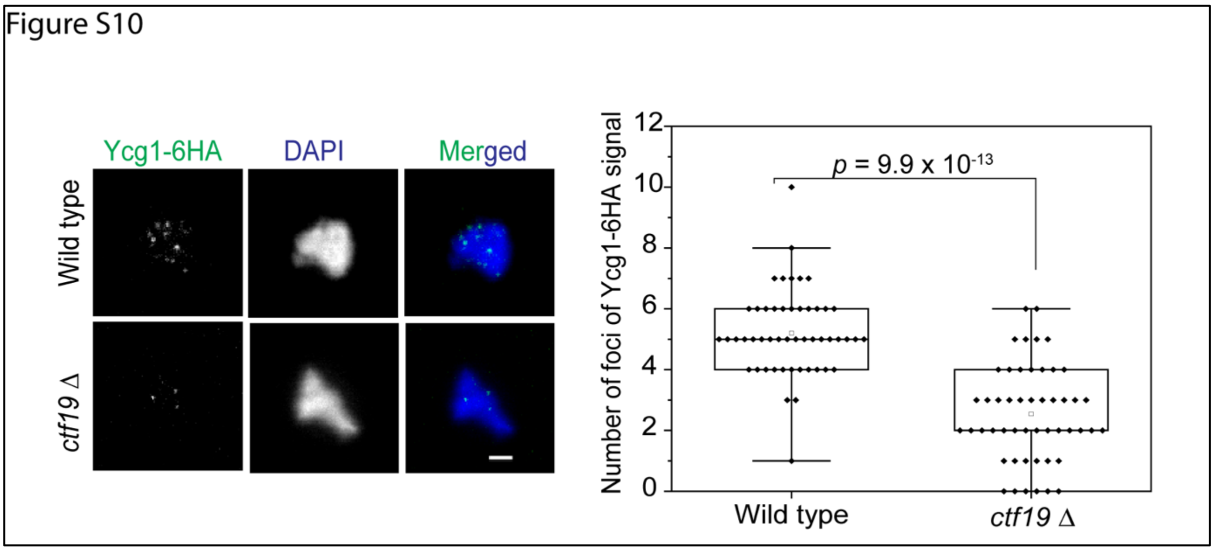

**Fig. S10. Reduced association of Ycg1 with chromatin in the absence of Ctf19 in meiosis.** Chromatin spreads showing chromatin-condensin association in the metaphase I arrested wild type (SGY9232) and *ctf19Δ* (SGY9233) cells harboring homozygous *YCG1-6HA* allele. Anti-HA antibodies (12CA5) were used to detect Ycg1-6HA. The left panel shows the representative images of the spreads, and the right panel represents the box plots showing the number of Ycg1-6HA foci per nuclear spread of the indicated strains. The statistical significance *p* value was estimated by the student's t-test for the mean. (N = 50 from two independent experiments, scale bar = 2 μm).

Figure S11:

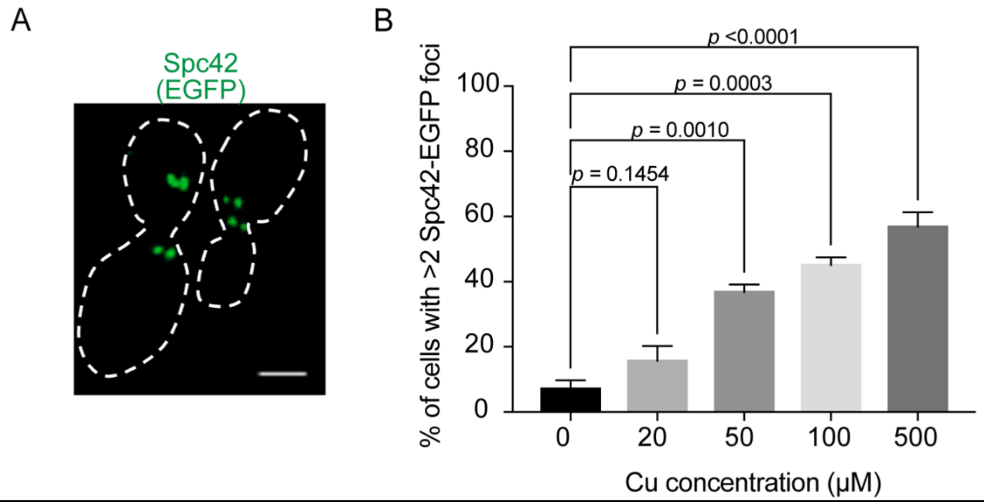

**Fig. S11. Overexpression of Cdc5 leads to supernumerary SPBs.** (A) Representative image showing multiple Spc42-EGFP foci in the wild type mitotic cells (SGY18008) harboring *CDC5* under *CUP1* promoter. (B) The percentage of these cells showing >2 SPB-EGFP foci when grown in YPD with indicated concentrations of copper. The statistical significance *p* value was estimated by the two-tailed student's t-test for the mean. (N = 50 from two independent experiments, scale bar = 2 µm).

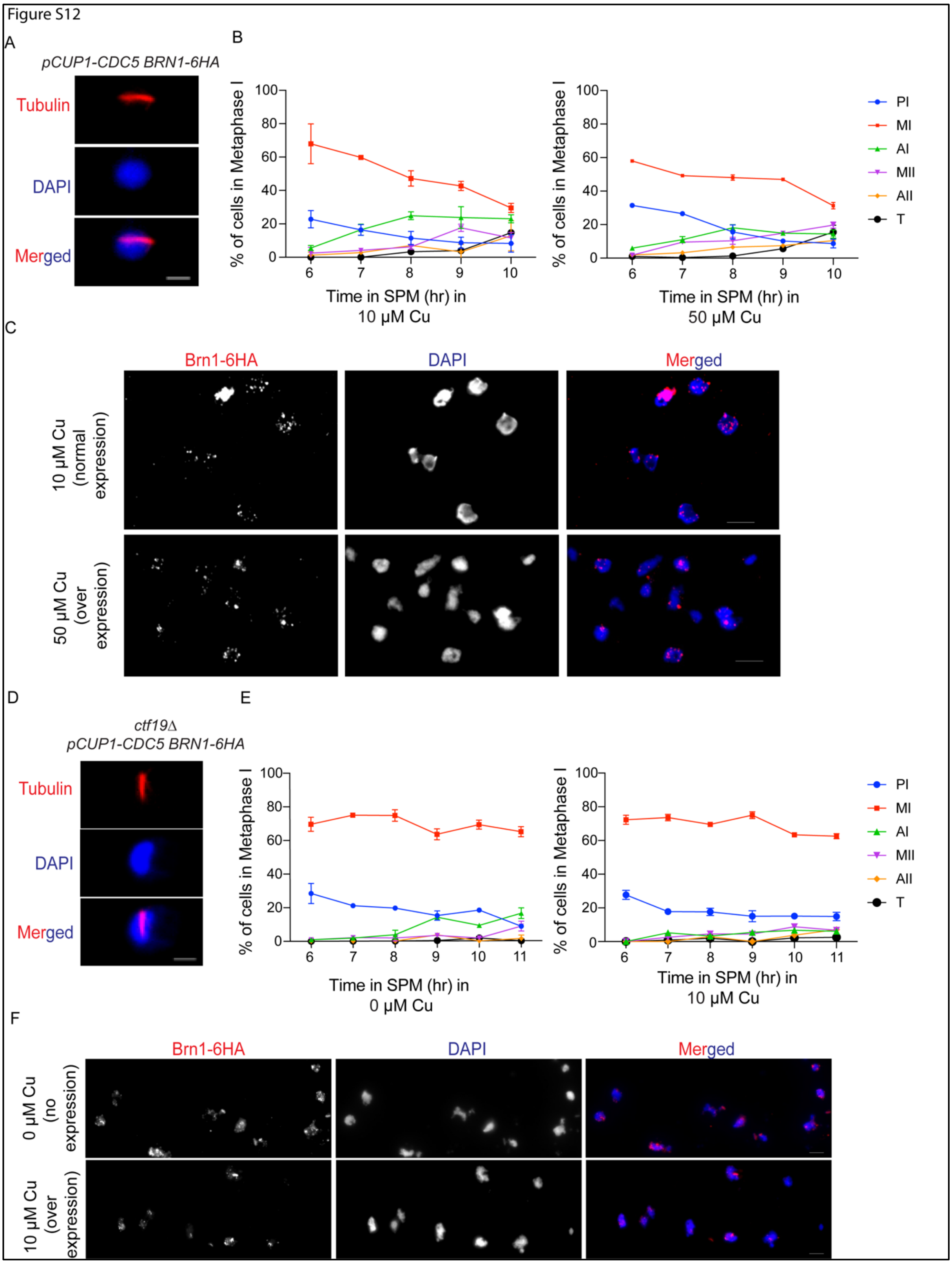

**Figure S12: Misregulation of Cdc5 influences chromatin-condensin association** (A) Representative images showing the indicated strain at metaphase I by immunofluorescence assay. (B) Bar graphs showing the percentage of metaphase I cells harboring *pCUP1-CDC5 BRN1-6HA*

(SGY18007) alleles at the indicated time points following their release into SPM supplemented with indicated concentrations of copper. (C) Field view of the chromatin spreads presented in Fig. 9C. (D) Same as in A. (E) Bar graphs showing the percentage of metaphase I cells harboring *ctf19Δ pCUP1-CDC5 BRN1-6HA* (SGY18015) alleles at the indicated time points following their release into SPM supplemented with indicated concentrations of copper. (F) Field view of the chromatin spreads presented in Fig. 9F. Scale bar = 5 μM.

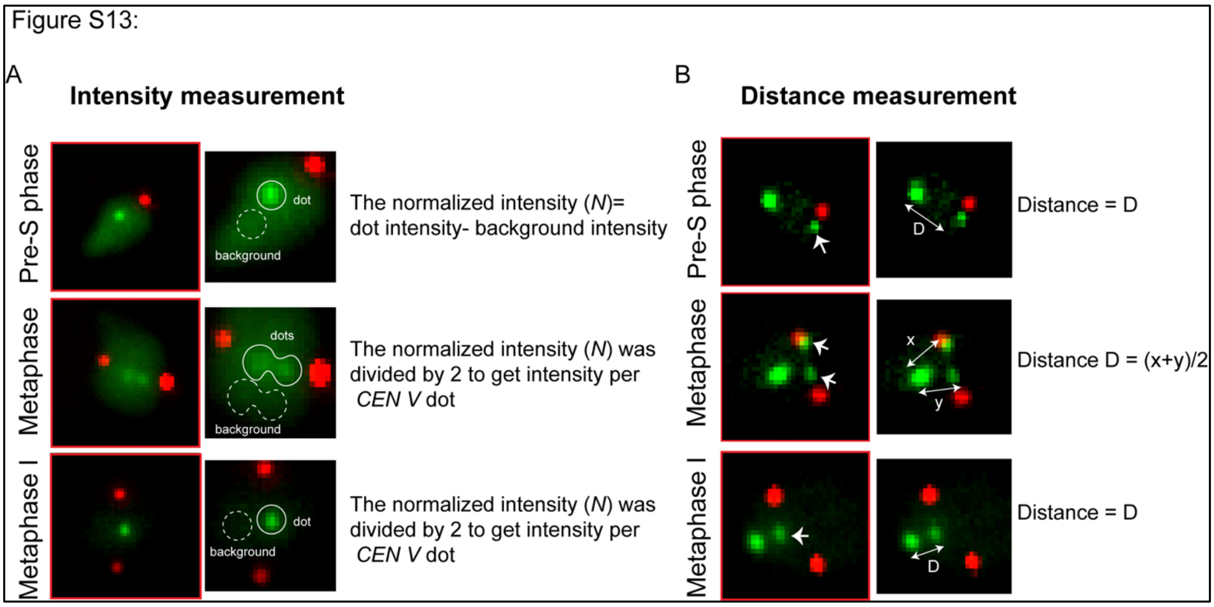

**Figure S13. The strategy adopted for measurement of intensity and distance in chromosome fluorescence-marked strains** (A) Fluorescence intensity of TetO/TetR-GFP as a read-out for chromatin compaction. Representative images of the cells in pre-S phase, metaphase and metaphase I, harboring $[TetO]_{224}$ array integrated 1.4 kb away from *CEN V* and expressing TetR-GFP (green) and SPB-mCherry (red). The normalized fluorescence intensity (*N*) of a focus was obtained by subtracting the background intensity from the total fluorescence intensity measured by ImageJ software. For metaphase and metaphase I cells of mitosis and meiosis, respectively, the *N* value was divided by 2 to get intensity per *CEN V* dot. (B) Distance between two fluorescently-marked loci of chromosome as a readout for axial contraction. Representative images of the cells in pre-S phase, metaphase and metaphase I, harboring $[TetO]_{224}$ and $[TetO]_{448}$ array inserted 1.4 kb and 21 kb away from *CEN V* and *TEL V*, respectively and expressing TetR-GFP (green) and SPB-mCherry (red). 3D distance between light (*CEN V*) and bright dot (*TEL V*) was measured using Imaris software. In metaphase cells, as there are two *CEN V* dots, the final distance (D) was calculated by taking an average of the distances between *TEL V* and both *CEN V* dots (x and y). The single-headed arrow indicates the *CEN V* dot.

### SUPPLEMENTAL TABLES

**Table S1: List of yeast strains used in this study**

| S.No. | Strain code | Genotype (all strains are of SK1 background) | Reference |
|---|---|---|---|
| 1 | SGY9012 | *MAT*a/α, *leu2::*TetR-*GFP::LEU2, CENV::tetOX224::HIS3, SPC42-mCherry::KanMx / SPC42-mCherry::KanMx* | This study |
| 2 | SGY9019 | *MAT*a/α, *leu2::*TetR-*GFP::LEU2, CENV::tetOX224::HIS3, SPC42-mCherry::KanMx , ctf19Δ::URA3/ SPC42-mCherry::KanMx, ctf19Δ::URA3* | This study |
| 3 | SGY9168 | *MAT*a/α, *leu2::*TetR-*GFP::LEU2, CENV::tetOX224::HIS3, SPC42-mCherry::KanMx , slk19Δ::URA3/ SPC42-mCherry::KanMx, slk19Δ::HPHMX* | This study |
| 4 | SGY9020 | *MAT*a/α, *cdc20::pCLB2-CDC20:: KanMx, slk19Δ::URA3 /cdc20::pCLB2-CDC20:: KanMx, NET1-GFP::TRP1, slk19Δ::URA3* | This study |
| 5 | SGY329 | *MAT*a/α, *cdc20::pCLB2-CDC20:: KanMx /cdc20::pCLB2-CDC20:: KanMx, NET1-GFP::TRP1* | Mehta et al., 2014 |

| | | | |
|---|---|---|---|
| 6 | SGY327 | *MAT*a/α, *cdc20::pCLB2-CDC20:: KanMx, ctf19Δ:: KanMx, REC8-6HA::HIS3/cdc20::pCLB2-CDC20:: KanMx, ctf19Δ:: KanMx, NET1-GFP::TRP1, REC8-6HA::HIS3* | Mehta et al., 2014 |
| 7 | SGY9092 | *MAT*a/α, *promURA3::*TetR*::GFP::LEU2,telV::tetOX448::URA3 (between BMH1 and PDA1), CENV::tetOX224::HIS3, SPC42-mCherry::KanMx/ SPC42-mCherry::KanMx* | This study |
| 8 | SGY9124 | *MAT*a/α, *promURA3::*TetR*::GFP::LEU2,telV::tetOX448::URA3 (between BMH1 and PDA1), CENV::tetOX224::HIS3, SPC42-mCherry::KanMx4, ctf19Δ::HPHMX / SPC42-mCherry::KanMx, ctf19Δ::URA3* | This study |
| 9 | SGY9025 | *MAT*a/α, *cdc20::pCLB2-CDC20:: KanMx, slk19Δ::URA3,BRN1-6HA::HIS3 /cdc20::pCLB2-CDC20:: KanMx, slk19Δ::URA3, BRN1-6HA::HIS3* | This study |
| 10 | SGY9163 | *MAT*a/α, *BRN1-6HA::HIS3, CDC20-6HA-AID:: KanMx, pADH-OsTIR1::URA3/ BRN1-6HA::HIS3, CDC20-6HA-AID:: KanMx, pADH-OsTIR1::URA3* | This study |
| 11 | SGY9165 | *MAT*a/α, *BRN1-6HA::HIS3, CDC20-6HA-AID:: KanMx, pADH-OsTIR1::URA3, ctf19Δ::HPHMX / BRN1-6HA::HIS3, CDC20-6HA-AID:: KanMx, pADH-OsTIR1::URA3, ctf19Δ::HPHMX* | This study |
| 12 | SGY283 | *MAT*a/α, *cdc20::pCLB2-CDC20:: KanMx, BRN1-6HA::HIS3 /cdc20::pCLB2-CDC20:: KanMx, BRN1-6HA::HIS3* | Mehta et al., 2014 |

| | | | |
|---|---|---|---|
| 13 | SGY284 | *MAT*a/α, *ctf19Δ::KanMx, cdc20::pCLB2-CDC20:: KanMx, BRN1-6HA::HIS3 / ctf19Δ::KanMx, cdc20::pCLB2-CDC20:: KanMx, BRN1-6HA::HIS3* | Mehta et al., 2014 |
| 14 | SGY916 9 | *MAT*a/α, *promURA3::*TetR*::GFP::LEU2,telV::tetOX448::URA3 (between BMH1 and PDA1), CENV::tetOX224::HIS3, SPC42-mCherry::KanMx4, DBF4-9MYC::HPHMX / SPC42-mCherry::KanMx, DBF4-9MYC::HPHMX* | This study |
| 15 | SGY918 9 | *MAT*a/α, *SGO1-EGFP::TRP1, NDC80-mCherry:: KanMx/ SGO1-EGFP::TRP1, NDC80-mCherry:: KanMx* | This study |
| 16 | SGY919 3 | *MAT*a/α, *SGO1-EGFP::TRP1, NDC80-mCherry:: KanMx, ctf19Δ::HPHMX / SGO1-EGFP::TRP1, NDC80-mCherry:: KanMx, ctf19Δ::HPHMX* | This study |
| 17 | SGY915 2 | *MAT*a/α, *cdc20::pCLB2-CDC20:: KanMx, CDC5-6HA::HIS3 /cdc20::pCLB2-CDC20:: KanMx, CDC5-6HA::HIS3* | This study |
| 18 | SGY914 8 | *MAT*a/α, *ctf19*Δ*:: KanMx , cdc20::pCLB2-CDC20:: KanMx, CDC5-6HA::HIS3 / ctf19*Δ*:: KanMx, cdc20::pCLB2-CDC20:: KanMx, CDC5-6HA::HIS3* | This study |
| 19 | SGY913 9 | *MAT*a/α, *CDC5-6HA::HIS3 / CDC5-6HA::HIS3* | This study |
| 20 | SGY914 0 | *MAT*a/α, *CDC5-6HA::HIS3, ctf19Δ::HPHMX / CDC5-6HA::HIS3, ctf19Δ::HPHMX* | This study |
| 21 | SGY923 2 | *MAT*a/α, *cdc20::pCLB2-CDC20:: KanMx, YCG1-6HA::HIS3 /cdc20::pCLB2-CDC20:: KanMx, YCG1-6HA::HIS3* | This study |

| | | | |
|---|---|---|---|
| 22 | SGY9233 | *MAT*a/α*, cdc20::pCLB2-CDC20:: KanMx, YCG1-6HA::HIS3, ctf19Δ::HPHMX /cdc20::pCLB2-CDC20:: KanMx, YCG1-6HA::HIS3, ctf19Δ::HPHMX* | This study |
| 23 | SGY9210 | *MAT*α*, CDC5-6HA::HIS3, CDC20-6HA-AID:: KanMx, pADH-OsTIR1::URA3* | This study |
| 24 | SGY9208 | *MAT*α*, CDC5-6HA::HIS3, CDC20-6HA-AID:: KanMx, pADH-OsTIR1::URA3, ctf19Δ::HPHMX* | This study |
| 25 | SGY7098 | *MATa/α, IME1-6HA::HIS3/ IME1-6HA::HIS3, CLN2-9MYC::KANMX/CLN2-9MYC::KANMX* | This study |
| 26 | SGY9194 | *MAT*a/α*, cdc20::pCLB2-CDC20:: KanMx, SGO1-9MYC::HPHMX /cdc20::pCLB2-CDC20:: KanMx, SGO1-9MYC::HPHMX* | This study |
| 27 | SGY9195 | *MAT*a/α*, cdc20::pCLB2-CDC20:: KanMx, SGO1-9MYC::HPHMX, ctf19Δ::KanMx /cdc20::pCLB2-CDC20:: KanMx, SGO1-9MYC::HPHMX, ctf19Δ::KanMx* | This study |
| 28 | SGY18007 | *MATa/α, cdc5::pCUP1-CDC5::KanMx, BRN1-6HA::HIS3 /cdc5::pCUP1-CDC5::KanMx, BRN1-6HA::HIS3* | This study |
| 29 | SGY18015 | *MATa/α, cdc5::pCUP1-CDC5::KanMx, BRN1-6HA::HIS3, ctf19Δ::HPHMX /cdc5::pCUP1-CDC5::KanMx, BRN1-6HA::HIS3, ctf19Δ::HPHMX* | This study |
| 30 | SGY9277 | *MAT*a/α*, cdc20::pCLB2-CDC20:: KanMx, ctf3Δ::URA3,BRN1-6HA::HIS3 /cdc20::pCLB2-CDC20:: KanMx, ctf3Δ::URA3, BRN1-6HA::HIS3* | This study |
| 31 | SGY9225 | *MAT*a/α*, cdc20::pCLB2-CDC20:: KanMx, ,IPL1-6HA::HIS3, , MTW1- 9MYC:: HPHMX /cdc20::pCLB2-CDC20:: KanMx,, IPL1-6HA::HIS3, , MTW1- 9MYC:: HPHMX* | This study |

| | | | |
|---|---|---|---|
| 32 | SGY9228 | *MAT*a/α, *ctf19Δ::LEU2 , cdc20::pCLB2-CDC20:: KanMx, ,IPL1-6HA::HIS3, , MTW1- 9MYC:: HPHMX / ctf19Δ::LEU2, cdc20::pCLB2-CDC20:: KanMx,, IPL1-6HA::HIS3, , MTW1- 9MYC:: HPHMX* | This study |
| 33 | SGY18008 | *MATa, cdc5::pCUP1-CDC5:: KanMx, SPC42-EGFP::TRP1 ,BRN1-6HA::HIS3* | This study |

**Table S2: Primer sequences used for ChIP experiments**

| | | |
|---|---|---|
| 1 | *CEN III* forward primer | GATCAGCGCCAAACAATATGG |
| 2 | *CEN III* reverse primer | AACTTCCACCAGTAAACGTTT |
| 3 | *CEN IV* forward primer | GCTTGCAAAAGGTCACATGC |
| 4 | *CEN IV* reverse primer | GAGCAGGTTTTATGTTTCGG |
| 5 | ChrIV pericentromere (P4) forward primer | CTTCTGGCTTGTTCACAACT |
| 6 | ChrIV pericentromere (P4) reverse primer | GGCAATTGTAGGTGGACTAA |
| 7 | ChrIV arm (A4) forward primer | GTCGATGGTTTCATTCAGAT |
| 8 | ChrIV arm (A4) reverse primer | CAATGGAGAGAGTGGATGTT |
| 9 | *TUB2* forward primer | CTTGTAGACAGCGTCATGG |
| 10 | *TUB2* reverse primer | CAGATGTCATAAAGTGCTTCG |
